# pLM-bVAE: protein language model-enhanced binary representation improves quantum annealing-based protein sequence design

**DOI:** 10.64898/2026.11.05.764447

**Authors:** Hiromi H Ueda, Andrejs Tučs, Koji Tsuda, Mitsuo Umetsu, Yutaka Saito

## Abstract

Quantum annealing (QA) has been explored as an approach for protein design in vast combinatorial sequence spaces. Applying QA to protein sequence design requires amino acid sequences to be encoded as binary vectors, and optimized vectors to be decoded back into amino acid sequences. Thus, both the quality of binary encoding and sequence reconstruction accuracy are important for effective QA-based optimization. Binary variational autoencoder (bVAE) has been used to map protein sequences to binary representations in QA-based protein sequence design. However, bVAE was originally developed for natural language modeling, and its suitability for proteins has not been systematically evaluated. Here, we present pLM-bVAE, which incorporates protein representations derived from a pretrained protein language model (pLM) into the original bVAE model. In computational experiments on three public protein variant datasets, pLM-bVAE showed higher sequence reconstruction accuracy and binary encoding performance than bVAE. Moreover, we incorporated pLM-bVAE into a QA-based protein sequence design framework. Across all three datasets, pLM-bVAE generated more high-scoring sequences with greater sequence dissimilarity from the training and validation data than bVAE. Our results demonstrate that pLM-bVAE provides an effective binary representation for improving QA-based protein sequence design.

**Key Points:**

● We developed pLM-bVAE, a protein language model-enhanced binary representation method for QA-based protein sequence design.
● Compared with the existing method bVAE, pLM-bVAE achieved higher sequence reconstruction accuracy and better preservation of sequence relationships in the binary latent space.
● Incorporating pLM-bVAE into a QA-based protein sequence design framework improved optimization performance.
● QA using pLM-bVAE generated more high-scoring sequences with greater sequence dissimilarity from the training and validation data, compared with bVAE.

## Introduction

Machine learning-guided protein sequence design aims to identify proteins with desired functions from vast combinatorial sequence spaces, which has been applied to the engineering of antibodies [1], peptides [2], enzymes [3], fluorescent proteins [4], and other biomolecules. Diverse computational approaches have been developed for this purpose, including Bayesian optimization [3, 5, 6], evolutionary algorithms [2, 7, 8], generative modeling [9–11], and quantum annealing (QA)-based approaches [12–16]. In this study, we focus on QA-based protein sequence design. In QA-based optimization, candidate solutions are represented by binary variables; therefore, applying QA to protein sequence design requires a representation method that maps amino acid sequences to binary vectors and converts optimized binary vectors back into amino acid sequences.

Previous studies have used binary variational autoencoder (bVAE) to bridge the sequence space and the binary space for QA-based protein sequence design [14, 15]. In this framework, the encoder maps amino acid sequences to binary latent vectors, whereas the decoder reconstructs amino acid sequences from QA-optimized binary vectors. The quality of this mapping is therefore critical: the binary vectors should preserve relationships among amino acid sequences, and the decoder should accurately recover amino acid sequences from the binary vectors. However, the existing bVAE architecture was originally developed for natural language modeling [17] and does not explicitly incorporate protein-specific sequence representations. This raises the possibility that representations better adapted to protein sequences could yield binary vectors that better preserve relationships among protein sequences while enabling more accurate sequence reconstruction.

Here we developed pLM-bVAE, which incorporates residue-level representations derived from a pretrained protein language model (pLM) into the bVAE model. We evaluated pLM-bVAE on three public protein variant datasets by assessing both sequence reconstruction accuracy and the extent to which relationships among protein sequences were preserved in the binary latent space. We further compared multiple pLMs and examined the effect of hyperparameter optimization on model performance. Finally, we incorporated the resulting pLM-bVAE into a QA-based protein sequence design framework and evaluated whether improved binary representation led to improved sequence optimization performance. Our results show that pLM-bVAE provides more accurate sequence reconstruction and binary vectors that better preserve sequence relationships than the conventional bVAE, leading to improved QA-based protein optimization performance across all three datasets.

## Materials and Methods

### Datasets and preprocessing

To train and evaluate pLM-bVAE, three public protein variant datasets (i.e., MHC-I binding peptides [18–20], Trastuzumab-scFv CDRH3 variants [21, 22] and AAV capsid surface region variants [23]) were used. The preprocessed datasets are summarized in Table 1.

**Table 1.** The preprocessed datasets used in this study.

| Dataset | Sequence length in variant regions (a.a.) | Experimentally measured value | No. training data | No. validation data | No. test data |
|---|---|---|---|---|---|
| MHC-I binding peptides [18] | 8–30 | log<sub>*e*</sub> *IC*<sub>50</sub>(nM) | 9,728 | 1,216 | 1,216 |
| Trastuzumab-scFv CDRH3 variants [21] | 14–18 | log<sub>10</sub> *K*<sub>*D*</sub>(nM) | 7,672 | 959 | 960 |
| AAV capsid surface region variants [23] | 28–43 | Viability score | 13,394 | 1,674 | 1,675 |

The original dataset of MHC-I binding peptides was downloaded from the “binding_data_2013.zip” available at https://tools.iedb.org/main/datasets/. This file contained data for multiple MHC-I alleles. To restrict the analysis to a single MHC-I target, peptides targeting the HLA-A*02:01 MHC-I molecule were extracted, as this allele is relatively frequent across multiple human populations [24] and a sufficiently large number of experimentally measured peptide IC_50_ (nM) was available for this allele. The IC_50_ values were natural-log-transformed and used as the scores of the peptide sequences. The size of the preprocessed dataset was 12,160.

The original dataset of Trastuzumab-scFV CDRH3 variants was downloaded from “YM_0989” in the “aalphabio/open-alphaseq” repository on Hugging Face using the Hugging Face Datasets library. This dataset included negative control data. The scFv CDRH3 variants targeting HER2 and annotated as “HER2-OPT_AGA2-POI” in “matalpha_description” were extracted. To remove less reliable measurements, variants annotated as True for both “above_background” and “sufficient_replicate_observations” were further extracted. Duplicate sequences were removed. The “alphaseq_affinity” values were experimentally measured by the AlphaSeq technique [21], which corresponds to log_10_ *K_D_* (nM). These values were used as the scores for the scFv CDRH3 variant sequences. The size of the preprocessed dataset was 9,591.

The original dataset of AAV capsid surface region variants was downloaded from the “library_w_selection_scores.csv.zip” at https://github.com/churchlab/Deep_diversification_AAV/tree/main/Data. Since this original dataset contained 243,481 variants, only AAV variants annotated as “rnn_designed_plus_rand_train_walked” in “category_x” were retained to reduce the dataset size for model training and evaluation. This subset contained highly diverse sequences with high viability scores [23] and was therefore suitable for model training. Following the same preprocessing method as the original study [23], variants with “GAS1_plasmid_N” values greater than 100 were retained. Variants with infinite “GAS1_virus_S” values were removed. Duplicate sequences were also removed, retaining the sequence entry with the largest “GAS1_virus_S” value. The “GAS1_virus_S” values represent the viability scores of AAV variants and were used as the scores for the AAV variant sequences. The size of the preprocessed dataset was 16,743.

### Model architecture

The architecture of bVAE was originally developed by [17]. The details of bVAE architecture and training procedure in our study are described in Supplementary data.

The pLM-bVAE architecture consists of a pLM encoder and a bVAE (Fig. 1A). The pLM encoders used in this study are listed in Table 2. The pretrained encoder models were obtained through the Huggingface API. The pLM encoder embedded full-length protein sequences into residue-level representation vectors. Then, the representation vectors corresponding to residues in the variant region were extracted from the full-length protein representations. The variant region was defined at the residue level for each dataset. In the MHC-I binding peptide dataset, the entire peptide sequence was treated as the variant region. For the CDRH3 variant dataset, the CDRH3 residues were defined according to the original studies [1, 21]. The AAV capsid surface region variant dataset contains the sequences corresponding to the 561—588 residues of the wild type AAV2 sequence (GenBank: YP_680426.1), which was treated as the variant region.

**Figure 1.**
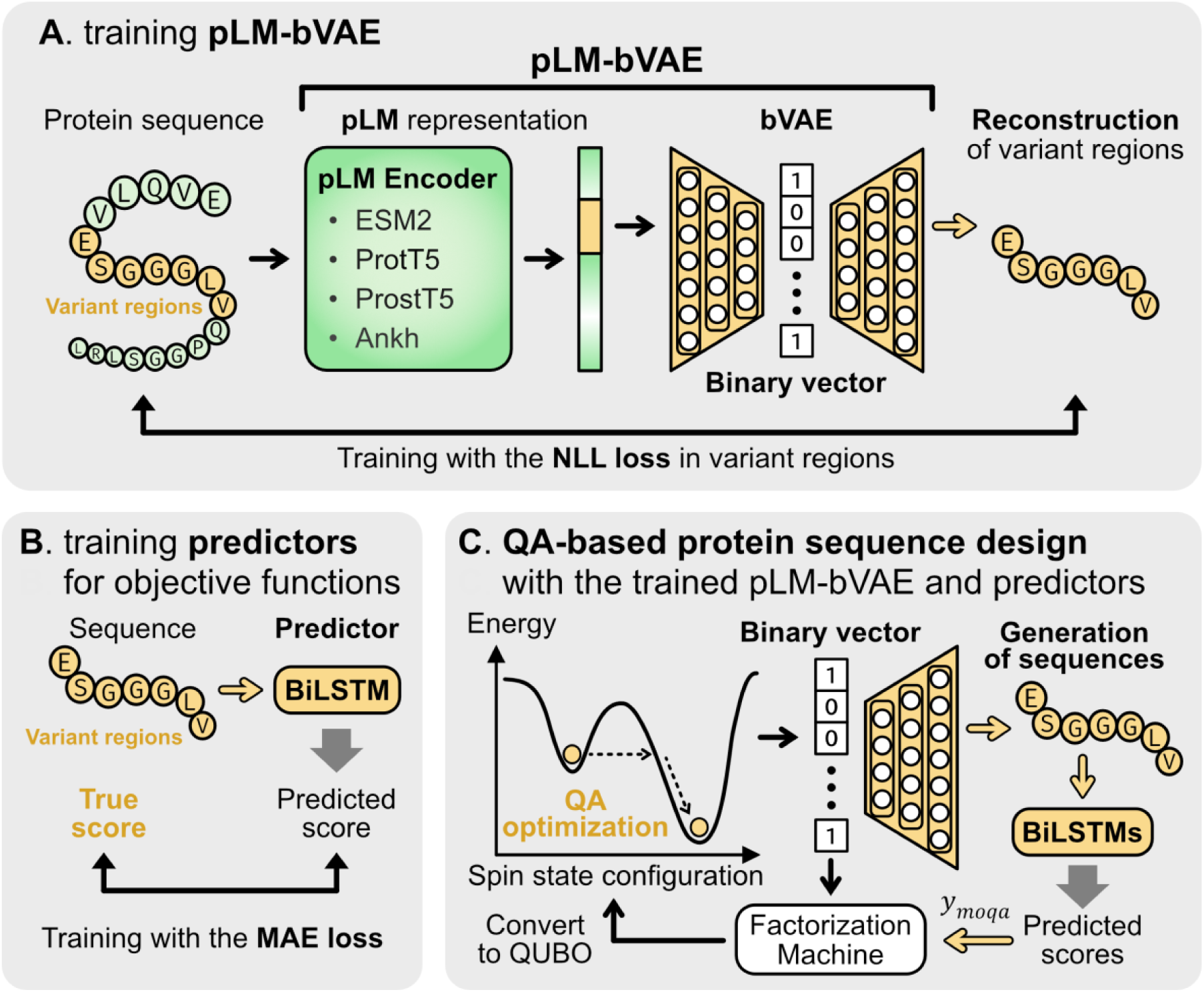
The overview of pLM-bVAE and workflow of QA-based protein sequence design. **(A)** The overview of training pLM-bVAE. We input a protein sequence to pLM-bVAE and obtain a binary vector that represents the variant regions of the protein sequence. The variant regions are defined by user at the residue level. pLM-bVAE is trained with the NLL loss in the variant regions. The parameters of bVAE component are trainable while the parameters of pLM encoder are frozen. **(B)** Training predictors for an objective function. BiLSTM predictors are trained with the MAE loss. **(C)** The workflow of QA-based protein sequence design. The aim of the workflow is to find high-scoring sequences by means of the trained predictors. Using the dataset of sequences and their predicted scores, *y_moqa_* values are calculated by the MOQA framework (Fig. S2). Using binary vectors corresponding to these sequences and *y_moqa_* values, FM is trained and subsequently converted to QUBO. QA optimization searches solutions for the QUBO, which samples binary vectors (spin state configurations) that minimize the corresponding Ising model energy. The sampled binary vectors are decoded into amino acid sequences using the trained pLM-bVAE decoder. In each iteration of QA optimization, sampled binary vectors are added to the current dataset and used to retrain FM to perform the next iteration.

**Table 2.** The pLM encoders applied in this study.

| Model | Number of parameters (encoder) | Encoder layers | Embedding size/residue | Huggingface model checkpoint |
|---|---|---|---|---|
| ESM2_8M [33] | 8 M | 6 | 320 | esm2_t6_8M_UR50D |
| ESM2_650M [33] | 650 M | 33 | 1280 | esm2_t33_650M_UR50D |
| ESM2_3B [33] | 3000 M | 36 | 2560 | esm2_t36_3B_UR50D |
| ProtT5 [34] | 1200 M | 24 | 1024 | prot_t5_xl_uniref50 |
| ProstT5 [35] | 1200 M | 24 | 1024 | ProstT5 |
| Ankh-base [36] | 736 M | 48 | 768 | ankh-base |
| Ankh-large [36] | 1900 M | 48 | 1536 | ankh-large |

The bVAE architecture was largely unchanged from the original architecture described above (Fig. S1). One difference was the embedding used by the decoder: while the bVAE decoder used the same amino acid descriptor as that used by the bVAE encoder for the embedding weights, the pLM-bVAE decoder used trainable embedding parameters with the same dimension as the bVAE for the embedding weights. During pLM-bVAE training, the reconstruction loss was defined as the sequence length-normalized negative log-likelihood (NLL) of the ground-truth sequences in the variant region:

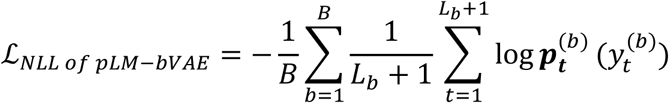

where *B* denotes the mini-batch size, *L_b_* denotes the length of the *b*-th amino acid sequence in the variant regions, and 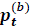 and 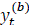 respectively denote the predicted probability distribution and its ground-truth token at the position *t*. The position *t* + 1 corresponds to the <eos> token at the terminus. During the inference, the most likely amino acid was selected from the predicted probability distribution at each decoding step. Sequence generation was terminated when the <eos> token was generated.

### Training and hyperparameter optimization procedure

The sequence data of each preprocessed dataset were randomly split into training, validation, and test sets at a ratio of 0.8: 0.1: 0.1 (Table 1). The bVAE and pLM-bVAE were trained on the same training set, validated on the same validation set, and tested on the same test set. The hyperparameters of pLM-bVAE were initially set to the same values as those used for bVAE in the previous study [15]: the number of encoder and decoder layers was 1, the optimizer was AdamW (weight_decay = 0), the hidden size was 256, the latent dimension was 64, the dropout rate was 0.2, the number of tokens per batch was 250, the number of epochs was 100, the learning rate was 0.001, the gradient clipping norm was 0.2, and the gradient accumulation step was 1.

To optimize the hyperparameter of pLM-bVAE, the Optuna package was employed [25]. Under the Optuna framework, Bayesian optimization based on the tree-structured Parzen estimator (TPE) sampler was performed for 200 trials. The searched hyperparameters were the number of encoder and decoder layers, the weight decay of AdamW, the hidden size, the dropout rate, the input dropout rate, the standard deviation of input noise, the label smoothing, the learning rate, the gradient clipping norm, whether to use the modified projection layer, and the gradient accumulation step while the number of epochs was fixed at 300. For each trial, pLM-bVAE was trained using the sampled hyperparameter set and evaluated on the validation set using the sequence length-normalized NLL scores:

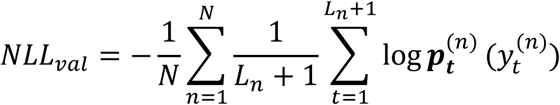

where *N* denotes the validation set size, *L_n_* denotes the length of the *n*-th amino acid sequence in the variant regions, and 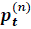 and 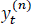 respectively denote the predicted probability distribution and its ground-truth token at the position *t*. The hyperparameter set that achieved the lowest validation NLL was selected as the best-performing set. Then, pLM-bVAE (HPO) was trained with the best hyperparameter set and the number of epochs showing the lowest *NLL_val_*.

### Evaluation of bVAE and pLM-bVAE

The trained bVAE, pLM-bVAE, and pLM-bVAE (HPO) were evaluated on the test set of each dataset. We designed the following evaluation measures in terms of sequence reconstruction accuracy, binary encoding performance, and total model performance.

The reconstruction accuracy was assessed using the following metrics. The ratio of sequences with a Levenshtein distance of 0 between the input and reconstructed sequences was given by:

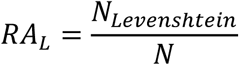

where *N_Levens_*_ℎ*tein*_ is the number of sequences with a Levenshtein distance of 0, and *N* is the test set size. Similarly, the ratio of sequences with a BLOSUM62 score between the input and reconstructed sequences greater than the 95th percentile of the BLOSUM62 scores obtained with bVAE was given by:

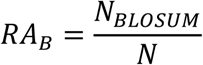

where *N_BLOSUM_* is the number of sequences with a BLOSUM62 score greater than this threshold, and *N* is the test set size. The BLOSUM62 score was calculated by pairwise global alignment using the BLOSUM62 substitution matrix, with gap-opening and gap-extension penalties of −10 and −0.5, respectively. PairwiseAligner in Biopython [26] was used for the alignment.

The binary encoding performance was assessed by measuring the preservation of sequence-distance relationships in the binary latent space. The Spearman correlation coefficient between the pairwise Levenshtein distances among test sequences and the pairwise Hamming distances among their corresponding binary vectors was given by:

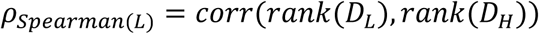

where *D_L_* and *D_H_* denote the pairwise Levenshtein distance and the pairwise Hamming distance, respectively. *rank*(·) denotes ranking with averaged ties, and *corr*(·) denotes the Pearson correlation coefficient. A higher positive correlation indicates better preservation of sequence-distance relationships in the binary latent space. Similarly, the Spearman correlation coefficient between the pairwise BLOSUM62 scores among test sequences and the pairwise Hamming distances among their corresponding binary vectors was given by:

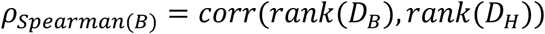

where *D_B_* and *D_H_* denote the pairwise BLOSUM62 score and the pairwise Hamming distance, respectively. Because higher BLOSUM62 scores indicate greater sequence similarity taking into account evolutionary divergence whereas larger Hamming distances indicate greater separation in the binary latent space, a stronger negative correlation indicates better preservation of evolutionary sequence relationships.

To compare overall model performance across the above metrics, the total model performance score was defined using rank-normalized measures of reconstruction accuracy and binary encoding performance. For each metric within each dataset, the models were ranked by their metric values. Let *r* denote the rank of a model, with *r* = 1 assigned to the worst-performing model and *r* = *M* assigned to the best-performing model, where *M* is the number of models. The rank-normalized reconstruction accuracy score was calculated as:

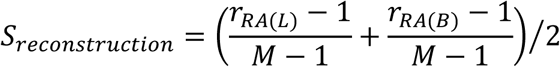

where *r_RA_*_(*L*)_ is the rank of the model for the *RA_L_* metric and *r_RA_*_(*B*)_ is the rank of the model for the *RA_B_* metric, respectively. Similarly, the rank-normalized binary encoding score was calculated as:

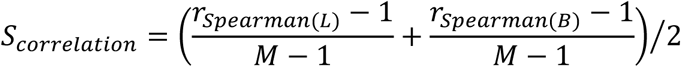

where *r_Spearman_*_(*L*)_ is the rank of the model for the *ρ_Spearman_*_(*L*)_ metric and *r_Spearman_*_(*B*)_ is the rank of the model for the *ρ_Spearman_*_(*B*)_ metric, respectively. The total model performance score for each dataset was calculated as:

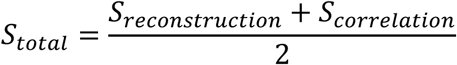

Overall model performance across the MHC-I binding peptide, CDRH3 variant, and AAV variant datasets was assessed by averaging the total model performance scores across the three datasets.

During the evaluation, generated sequences longer than 100 amino acids were considered aberrant decoder outputs and excluded from the evaluation.

### Training and evaluation of predictors for objective functions

To predict objective function values for variant region sequences generated by QA-based protein sequence design, predictors were trained with a regression task. The training, validation, and test splits were identical to those used for bVAE and pLM-bVAE described above (Table 1). These data consisted of pairs of variant region sequences and their corresponding scores. To train the predictors on the training and validation sets, the PARROT package was employed [27]. This package provides the framework for training bidirectional recurrent neural network based on long short-term memory cells (BiLSTM) [28] with the mean absolute error (MAE) loss function:

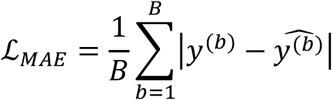

where *B* denotes the mini-batch size, which was set to 16, *y*^(*b*)^ is the score of the *b*-th data, and 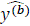 is its score predicted by the BiLSTM predictor. The detailed procedures for training and Bayesian hyperparameter optimization are provided in Supplementary data. For each dataset, we trained three BiLSTM predictors and evaluated the ensemble of them on the test set using the Pearson and Spearman correlation coefficients between the experimentally measured scores and the average of predicted scores from the three predictors.

For QA-based protein sequence design, predicted scores from the three predictors were separately used for non-dominated sorting in the multi-objective optimization by quantum annealing (MOQA) framework (Fig. S2). The average of predicted score from the three predictors was used as a computational oracle score to evaluate the optimization performance.

### QA-based protein sequence design framework combining bVAE/pLM-bVAE and predictors for objective functions

For QA-based protein sequence design, the MOQA framework [14, 15] was used in combination with the trained bVAE/pLM-bVAE and the trained predictors for the objective function described above (Fig. S2). This framework used factorization machine (FM), formulated as follows [29, 30]:

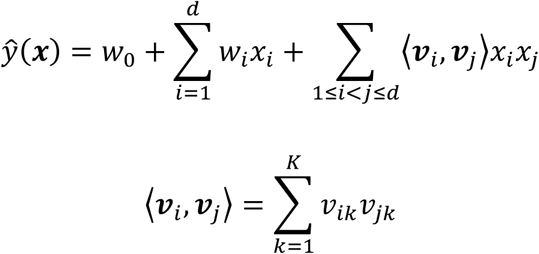

where ***x*** is a binary vector, and *d* is the dimension of the binary vector, which was set to 64. *w*_0_, *w_i_*, ndd *v_ik_* are the model parameters, and *K* is a hyperparameter that determines the dimensionality of the model parameters, which was set to 8. The FM was used as the surrogate model for the function of MOQA score, which evaluates generated sequences based on their ranks obtained by non-dominated sorting [31]. The trained FM was directly converted to a quadratic unconstrained binary optimization (QUBO) formulation and subsequently to an Ising model defined over spin-state configurations [30]. The Ising model was embedded into a quantum annealer, D-Wave Advantage System 6 [13]. Then, QA-based optimization was performed to search for spin state configurations that minimize the energy of the Ising model. Because the quantum annealer minimizes the QUBO, the predicted objective function scores were transformed as necessary so that better solutions corresponded to lower objective values during QA optimization.

The detailed procedure of the QA-based protein sequence design using the MOQA framework is described in Supplementary data.

### Evaluation of the QA-based protein sequence design

To extract unique sequences generated by QA-based sequence design, duplicate sequences and sequences present in the training and validation data (Train+Val data) were removed. To evaluate the performance of QA-based sequence design using bVAE or pLM-bVAE (HPO), the number of unique generated sequences with oracle scores higher than the 10-th highest oracle score in the Train+Val data was counted. This metric indicates the ability to generate sequences with high predicted objective scores. Furthermore, the average nearest-neighbor Levenshtein distance from the Train+Val data was calculated for generated sequences with oracle scores higher than this threshold. This metric quantifies the dissimilarity (novelty) of high-scoring generated sequences relative to the Train+Val data.

### Analysis of the QA-based optimization process in the binary latent space

To visualize the search process of QA-based optimization using bVAE/pLM-bVAE (HPO) in two dimensions, uniform manifold approximation and projection (UMAP) was applied to the 64-dimensional binary vectors of the initial data and those sampled during the QA-based optimization. To quantify the difference between the distributions of the randomly sampled binary vectors in the initial data and the binary vectors sampled by the QA-based optimization, maximum mean discrepancy (MMD) with a radial basis function (RBF) kernel was used [32]. The detailed procedure for calculating MMD is described in Supplementary data. Since the QA-based optimization was performed 10 times using different initial datasets, the values of MMD^2^ obtained from 10 runs were averaged. The averaged MMD^2^ quantifies the distributional difference between the initial binary vectors and those sampled during QA-based optimization.

To trace changes in the best oracle score over iterations of QA-based optimization, the Δ best oracle score was calculated by subtracting the best oracle score of the initial data from the best oracle score obtained up to each iteration.

## Results and Discussion

### Model overview and workflow of QA-based protein sequence design

In this study, we developed pLM-bVAE, a binary representation method for QA-based protein sequence design. The model consists of a pLM encoder and a bVAE (Fig. 1A). The pLM encoder embeds protein sequences into evolutionary-informed representations, which are then provided to the bVAE model for binary encoding and reconstruction. This architecture enables the mapping of protein sequences to binary latent vectors and their reconstruction back to amino acid sequences (Fig. S1). The pLM-bVAE was then applied to QA-based protein sequence design.

As illustrated in Fig. 1, the workflow of the QA-based protein sequence design consists of three components: training pLM-bVAE (Fig. 1A), training predictors for the objective functions (Fig. 1B), and QA-based protein sequence design with the trained pLM-bVAE and the trained predictors (Fig. 1C).

Training pLM-bVAE requires a protein variant sequence dataset (Fig. 1A). We preprocessed three public protein variant datasets listed in Table 1 (i.e., MHC-I binding peptides [18], Trastuzumab-scFv CDRH3 variants [21] and AAV capsid surface region variants [23]), and trained pLM-bVAE using the training and validation sets with the NLL loss. The variant regions of full-length sequences were defined at the residue level for each dataset (Materials and Methods), and used for binary encoding. Various types of pLM encoders are available (Table 2), but we primarily used ESM2_650M [33] which showed consistently high performance across the three datasets (Fig. 3). The parameters of the pLM encoder were frozen while the bVAE parameters were trainable.

The predictors for the objective functions were trained using the same training and validation sets as pLM-bVAE (Fig. 1B). The data consisted of pairs of variant sequences and their corresponding scores. BiLSTM predictors were trained using the training and validation sets with the MAE loss [27].

Using the trained pLM-bVAE decoder and the trained BiLSTM predictors, QA-based protein sequence design was performed for each dataset (Fig. 1C). First, we input 2,000 randomly sampled binary vectors to the decoder as the initial data. The decoder reconstructed 2,000 amino acid sequences, and the trained predictors output predicted scores from the sequences. Then, *y_moqa_* values were calculated under the MOQA framework (Fig. S2). Using the data of 2,000 pairs of the binary vectors and the *y_moqa_* values, FM was trained and subsequently converted to QUBO. The QUBO problem was solved using a D-Wave quantum annealer, which sampled 10 optimized binary vectors. These binary vectors were added to the existing dataset and used to retrain the FM. We repeated this QA process for 200 iterations, generating 2,000 optimized sequences. The entire procedure was repeated 10 times with different initial datasets, yielding approximately 20,000 optimized sequences in total.

The detailed procedures of the training bVAE/pLM-bVAE, the training predictors for objective functions, and the QA-based protein sequence design are given in Materials and Methods.

### pLM-bVAE improved reconstruction accuracy and binary encoding performance

To evaluate the model performance of bVAE/pLM-bVAE, we assessed reconstruction accuracy using Levenshtein distance between the input and output amino acid sequences using the trained bVAE/pLM-bVAE on the test data. The input sequences were encoded as 64-dimensional binary latent vectors by the trained bVAE/pLM-bVAE encoder, which in turn were decoded as the output sequences by the trained bVAE/pLM-bVAE decoder. The pLM encoder of pLM-bVAE was fixed to ESM2_650M to compare the performance of bVAE/pLM-bVAE, excluding the effect of different pLM encoder models. On the MHC-I binding peptide dataset (Fig. 2A), while the sequences reconstructed by the trained bVAE showed large Levenshtein distance, the trained pLM-bVAE reconstructed sequences with small Levenshtein distance from the input sequences (Fig. 2B). In addition, to estimate the reconstruction performance taking into account the similarity of amino acid residues, we calculated BLOSUM62 scores between the input and output sequences. Similarly to the assessment using Levenshtein distance, the trained pLM-bVAE showed higher BLOSUM62 scores than the trained bVAE (Fig. 2B). On other datasets (i.e., the Trastuzumab-scFv CDRH3 variants dataset and AAV capsid surface region variants dataset) (Fig. S3A and S4A), the trained pLM-bVAE also showed smaller Levenshtein distance and higher BLOSUM 62 scores than the trained bVAE (Fig. S3B and S4B). These results indicate that the ESM2-derived protein representations improved reconstruction accuracy, as assessed by Levenshtein distance and BLOSUM62 score. We further assessed the reconstruction performance of pLM-bVAE (HPO) that was trained with the Bayesian-optimized hyperparameters [25]. The trained pLM-bVAE (HPO) showed almost all reconstructed sequences with Levenshtein distance of 0 on the peptide dataset (Fig. 2B), meaning that almost all output sequences through the trained pLM-bVAE (HPO) exactly matched their input sequences. BLOSUM62 scores of the trained pLM-bVAE (HPO) were substantially higher than pLM-bVAE as well as bVAE. The similar tendencies were observed on the CDRH3 variant and AAV variant datasets (Fig. S3B and S4B). These results indicate that hyperparameter optimization further improved the reconstruction performance of pLM-bVAE.

**Figure 2.**
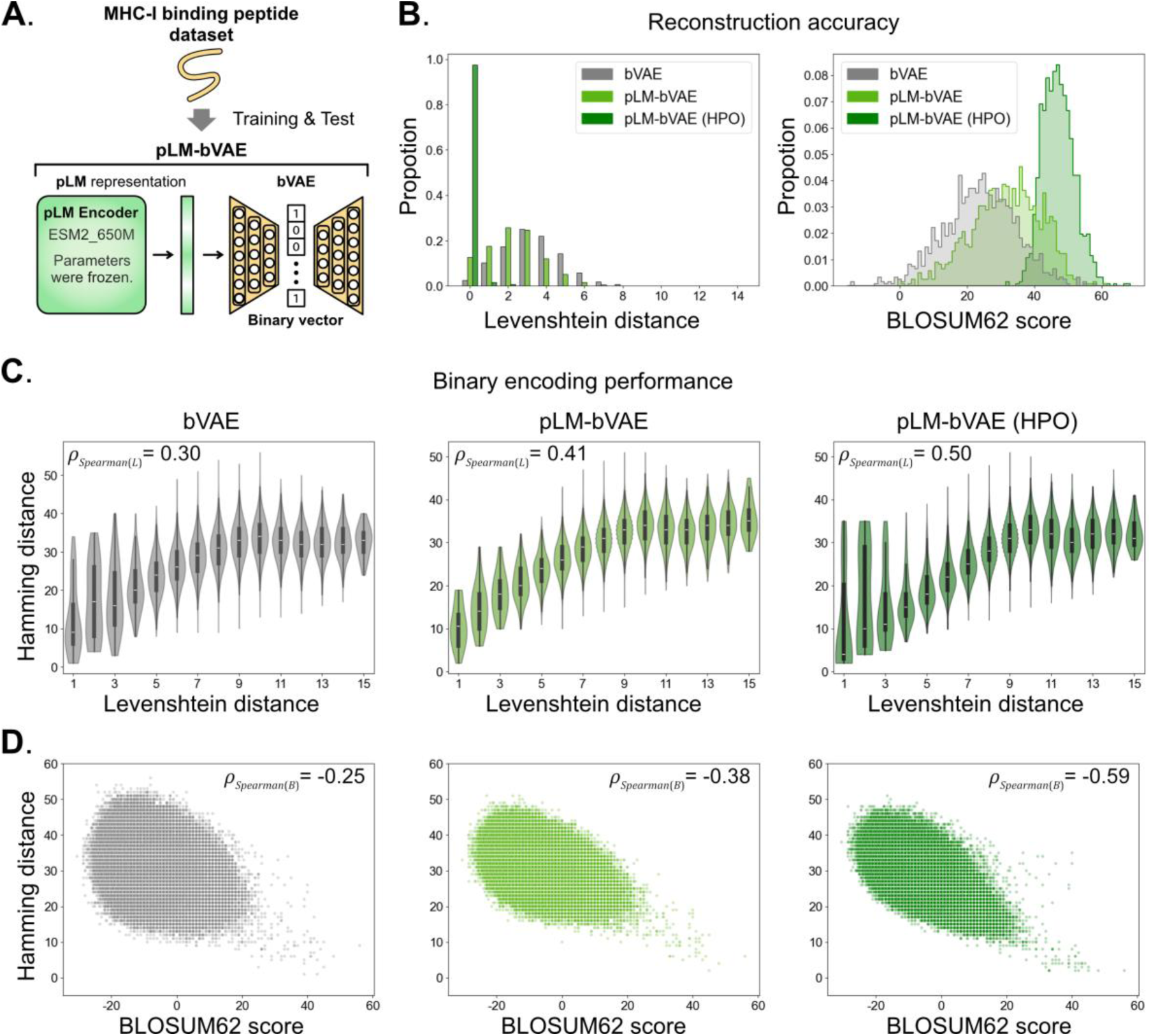
pLM-bVAE showed higher reconstruction accuracy and binary encoding performance than bVAE on the MHC-I binding peptide dataset. **(A)** To train and test bVAE and pLM-bVAE, the MHC-I binding peptide dataset was used. pLM-bVAE was composed of ESM2 650M encoder with the model parameters frozen and bVAE. The results on the scFv CDRH3 variant dataset and the AAV capsid surface region variant dataset are provided in Supplementary data. **(B)** Reconstruction accuracy of the trained bVAE, pLM-bVAE, and pLM-bVAE (HPO) on the test data. **Left**: Distributions of Levenshtein distance between input sequences and output sequences through the models. **Right**: Distributions of BLOSUM62 score between input sequences and output sequences through the models. **(C, D)** Binary encoding performance of the trained bVAE (left), pLM-bVAE (middle), and pLM-bVAE (HPO) (right) on the test data. **(C)** Correlation between pairwise Levenshtein distance of the test sequences and pairwise Hamming distance of the binary vectors encoded from their sequences. *ρ_Spearman_*_(*L*)_ shows the Spearman correlation coefficient value. **(D)** Correlation between pairwise BLOSUM62 score of the test sequences and pairwise Hamming distance of the binary vectors encoded from their sequences. *ρ_Spearman_*_(*B*)_ shows the Spearman correlation coefficient value.

**Figure 3.**
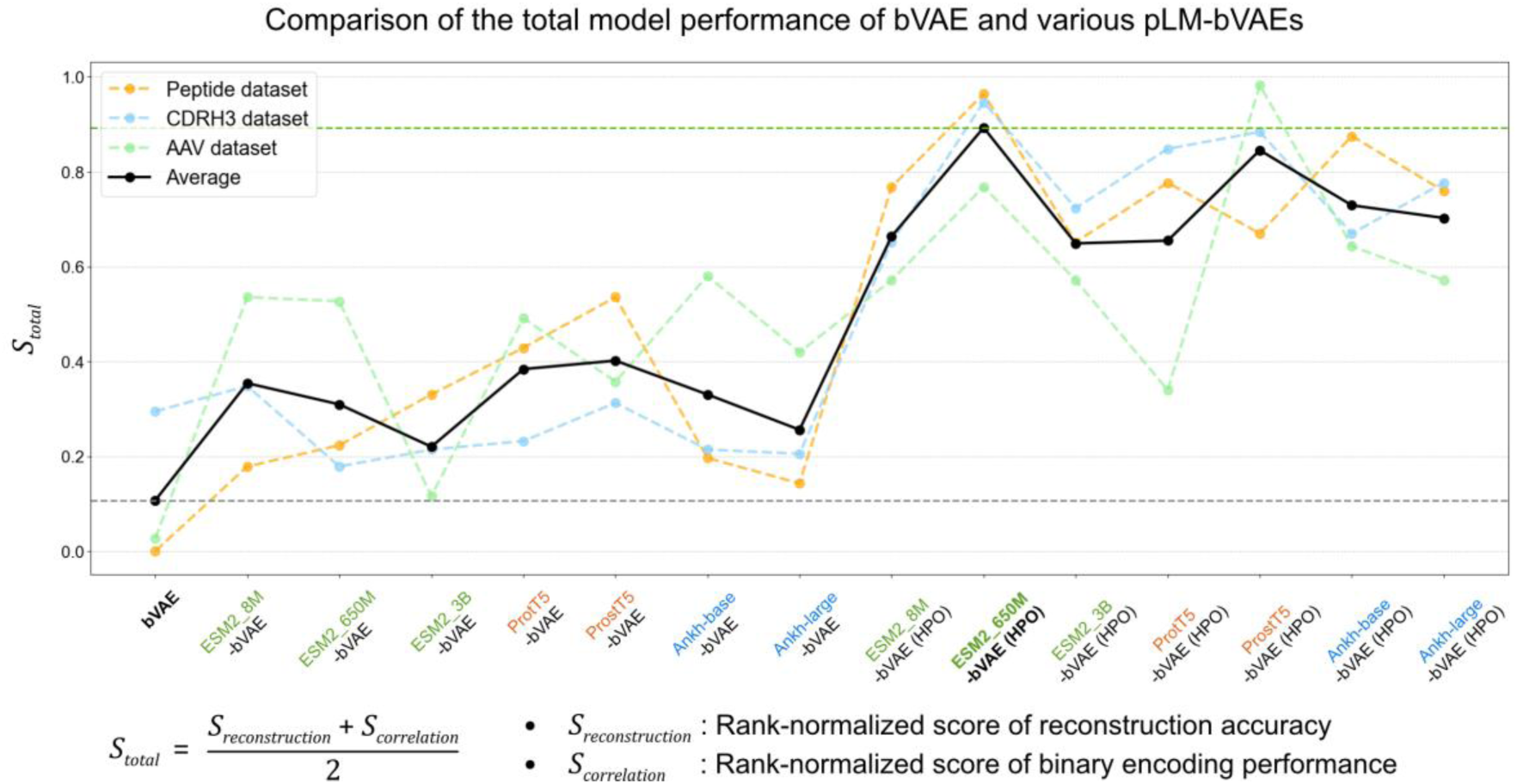
Comparison of the total model performance of bVAE and various pLM-bVAEs. *S_total_* was calculated by the aggregation of *S_reconstruction_* and *S_correlation_* that are the rank-normalized scores of reconstruction accuracy and binary encoding performance, respectively (Material and Methods). The dashed lines indicate *S_total_* on each dataset. The black line indicates averaged *S_total_* across the datasets. The horizontal gray and green dashed lines show the averaged *S_total_* values of bVAE and ESM2_650M-bVAE (HPO), respectively. The types of encoders are listed in Table 2.

Next, to evaluate binary encoding performance of bVAE/pLM-bVAE, we calculated the Spearman correlation coefficient between pairwise Levenshtein distance of the test sequences and pairwise Hamming distance of binary vectors encoded from these sequences. The high correlation between the pairwise distances of the sequences and their binary vectors means that the trained encoder preserves the distance information of the sequences in their binary latent vectors. We found that the trained pLM-bVAE/pLM-bVAE (HPO) outperformed the trained bVAE on all three datasets (Fig. 2C, S3C, S4C). The similar tendencies were observed in the correlation between pairwise BLOSUM62 score of the test sequences and pairwise Hamming distance of their binary vectors (Fig. 2D, S3D, S4D). These findings imply that the pLM representation in pLM-bVAE improved the preservation of sequence relationships captured by Levenshtein distance and BLOSUM62 similarity in the binary latent space.

Collectively, the proposed pLM-bVAE performed better than the original bVAE. Bayesian optimization of the hyperparameters further improved the performance of pLM-bVAE.

### Comparison of the total model performance of bVAE and various pLM-bVAEs

While we used ESM2_650M as the pLM encoder of pLM-bVAE (ESM2_650M-bVAE), there are various types of pLM encoders, such as the ESM2 with different model parameter sizes [33], encoder-decoder pLMs [34], pLMs that learn from the sequence and structural information [35], and protein-specific efficient pretraining pLMs [36]. It is possible that the combination of these pLM encoders with bVAE are more effective than ESM2_650M-bVAE. To examine this possibility, we compared the total model performance of bVAE/ESM2_650M-bVAE with other pLM encoders-combined bVAE on the same training, validation, and test data (Fig. 3). The total model performance score *S_total_* takes into account both reconstruction accuracy and binary encoding performance. The *S_total_* values were calculated by the aggregation of *S_reconstruction_* and *S_correlation_* that are the rank-normalized scores of reconstruction accuracy and binary encoding performance, respectively. *S_reconstruction_* are derived from the metrics of reconstruction accuracy, *RA_L_* (Fig. S5A, S6A, S7A) and *RA_B_* (Fig. S5B, S6B, S7B), and *S_correlation_* are derived from the metrics of binary encoding performance, *ρ_Spearman_*_(*L*)_ (Fig. S5C, S6C, S7C) and *ρ_Spearman_*_(*B*)_ (Fig. S5D, S6D, S7D). These metrics on each dataset are listed in Table S1, S2, and S3. The detailed calculation method is given in Materials and Methods. As shown by the averaged *S_total_* values in Fig. 3, while the trained bVAE was the worst-performing model, the trained ESM2_650M-bVAE (HPO) was the best-performing model across the datasets, suggesting that the ESM2_650M encoder is a high-performance pLM encoder for pLM-bVAE. The comparison of the total performance of pLM-bVAEs (HPO) using ESM2 with different model parameter sizes showed that ESM2_650M-bVAE (HPO) outperformed ESM2_3B-bVAE (HPO) that has larger model parameter size (Fig. 3). The same tendency was observed in the comparison of Ankh-base-bVAE (HPO) and Ankh-large-bVAE (HPO) that are the small and large parameter size models, respectively (Fig. 3). These results suggest that increasing the model parameter size does not necessarily improve the total model performance. The strong performance of ESM2_650M-bVAE (HPO) is consistent with previous studies reporting competitive transfer-learning performance of medium-sized ESM2 models when task-specific training data are limited [37, 38]. Based on the results, we decided to incorporate the trained ESM2_650M-bVAE (HPO) as pLM-bVAE (HPO) into the QA-based protein sequence design framework [30].

### pLM-bVAE improved QA-based generation of high-scoring sequences dissimilar to the Train+Val data

We examined the performance of bVAE/pLM-bVAE in QA-based protein sequence design (Fig. 4A). We used a previously-described QA-based design framework [14, 30] where we incorporated the trained bVAE or pLM-bVAE (HPO) as a binary representation method. Since this framework requires predictors for objective functions (Fig. S8A), we trained and evaluated the BiLSTM predictors [27] using the same training, validation, and test data as those used for bVAE/pLM-bVAE (HPO) (Fig. S8B). The three trained predictors output three predicted scores for each variant region sequence. These predicted scores were used as objective functions for non-dominated sorting in the MOQA framework [14, 15] (Fig. S2). The average of the three predicted score was used as the computational oracle score for the evaluation of QA-based protein sequence design performance. To evaluate the prediction accuracy of the ensemble of the three trained predictors, we calculated the Pearson and Spearman correlation coefficients between the experimentally measured scores and the averaged predicted scores, showing high correlations (Fig. S8C). These results support the use of the averaged predicted scores as computational oracle scores for comparing QA-based protein design performance.

**Figure 4.**
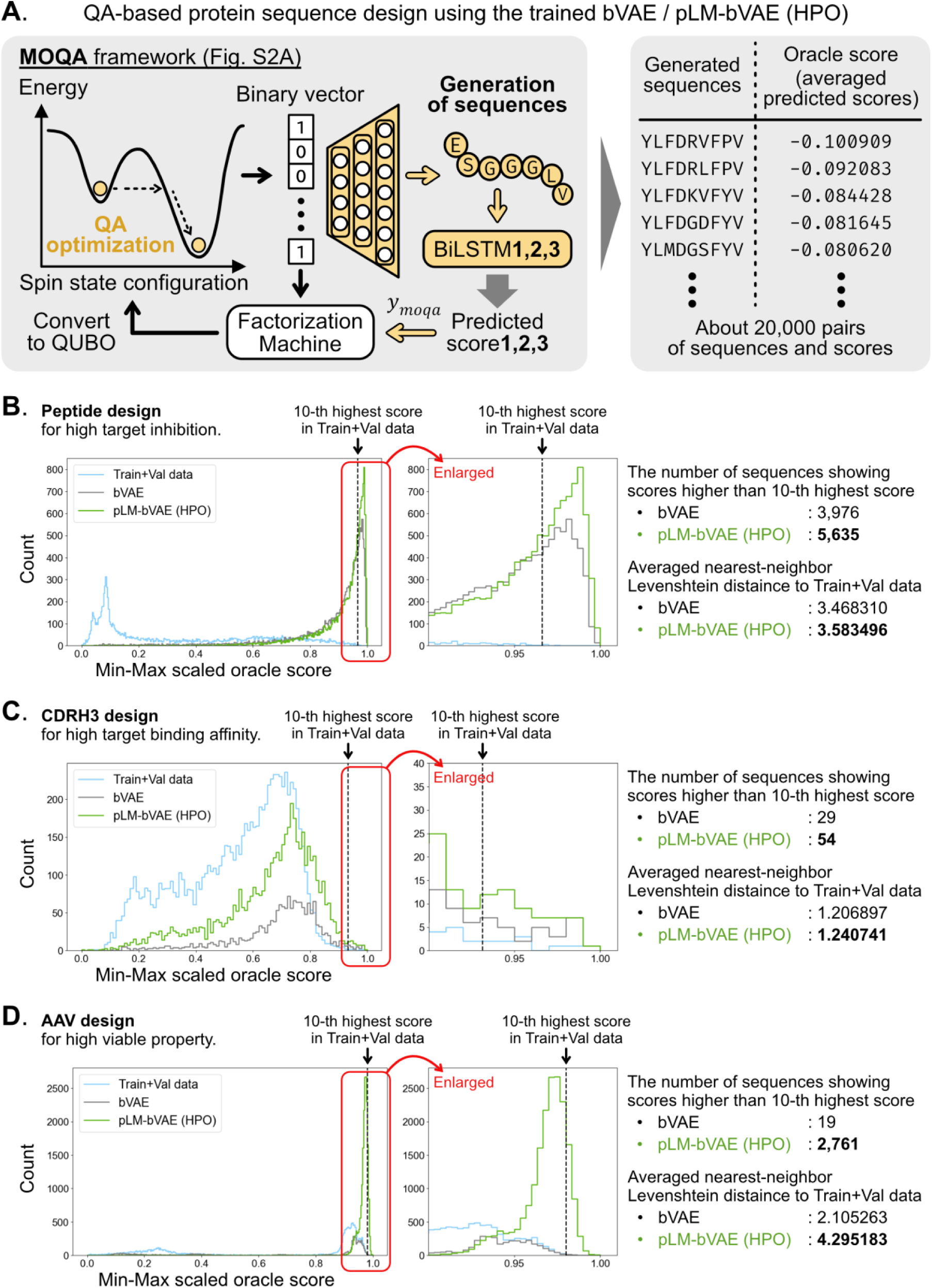
QA-based protein sequence design using pLM-bVAE generated a larger number of high-scoring sequences dissimilar to the Train+Val data. **(A)** Generation of protein sequences by QA-based sequence design using the trained bVAE/pLM-bVAE (HPO) under the MOQA framework (Fig. S2). Consequently, about 20,000 pairs of generated sequences and their computational oracle scores are obtained for each protein variant dataset. **(B, C, D) Left**: Distributions of the oracle scores are compared for the Train+Val data (used for training and validation of bVAE/pLM-bVAE (HPO) and the BiLSTM predictors), sequences generated by QA using bVAE, and sequences generated by QA using pLM-bVAE (HPO) on the peptide design **(B)**, the CDRH3 variant design **(C)**, and the AAV variant design **(D)**. The oracle score on the x-axis is Min-Max scaled. The vertical dashed line indicates the 10-th highest oracle score in the Train+Val data. **Middle**: The area near the 10-th highest score is enlarged. **Right**: (upper) The number of sequences showing oracle scores higher than the 10-th highest score. (lower) Averaged nearest-neighbor Levenshtein distance to the Train+Val data in the sequences with scores higher than the threshold.

Using the trained predictors and bVAE/pLM-bVAE (HPO) decoders, we performed the QA-based protein sequence design for each dataset. The detailed procedure of the QA-based protein sequence design is given in Materials and Methods and Supplementary data. For each dataset, about 20,000 optimized sequences were generated and sorted by the oracle score after duplicate sequences were removed (Fig. 4A). To evaluate the QA-driven design performance, we counted the number of generated sequences showing high oracle scores and assessed their sequence dissimilarity from the Train+Val data (Fig. 4B, 4C, 4D). A high oracle score was defined as a score higher than the 10-th highest oracle score in the Train+Val data for each dataset. Sequence dissimilarity from the Train+Val data was measured as the average nearest-neighbor Levenshtein distance. QA using pLM-bVAE (HPO) generated more sequences with high oracle scores and greater sequence dissimilarity from the Train+Val data than QA using bVAE across all three datasets (Fig. 4B, 4C, 4D). These results indicate that the reconstruction accuracy and binary encoding performance of bVAE/pLM-bVAE are associated with QA-based protein design performance.

### QA optimization process using pLM-bVAE (HPO) showed a more directed search pattern associated with improved QA optimization efficiency

It is possible that incorporating pLM-bVAE (HPO) alters the process of QA optimization in the binary latent space, thereby contributing to improved QA optimization efficiency and QA-based design performance. To investigate the mechanism by which pLM-bVAE improved the QA-based design performance, we visualized the QA optimization processes in the binary latent space for each dataset by applying UMAP to the binary vectors sampled by the QA searches (Fig. 5A, 5C, 5E). The QA searches started from the initial state obtained by training the FM on 2,000 randomly sampled binary vectors, and sampling of optimized binary vectors by QA was repeated for 200 iterations. We performed this QA optimization process 10 times using different initial data. Thus, as shown in Fig. 5A, 5C, 5E, while the randomly sampled binary vectors in the initial data were broadly distributed, the plots of binary vectors sampled by the QA searches formed trajectories over the iterations. As the iterations progressed, the sampled binary vectors progressively deviated from the initial distribution, indicating a directed shift in the search process. Moreover, the QA searches using pLM-bVAE (HPO) appeared to show a more pronounced shift than those using bVAE. To quantify this difference, we calculated the MMD between the distributions of the initial data and the binary vectors sampled by the QA searches using bVAE or pLM-bVAE (HPO) (Fig. 5A, 5C, 5E). The averaged MMD^2^values were larger for pLM-bVAE (HPO) than for bVAE across all datasets, indicating a greater distributional shift from the initial binary vector distribution during QA optimization.

**Figure 5.**
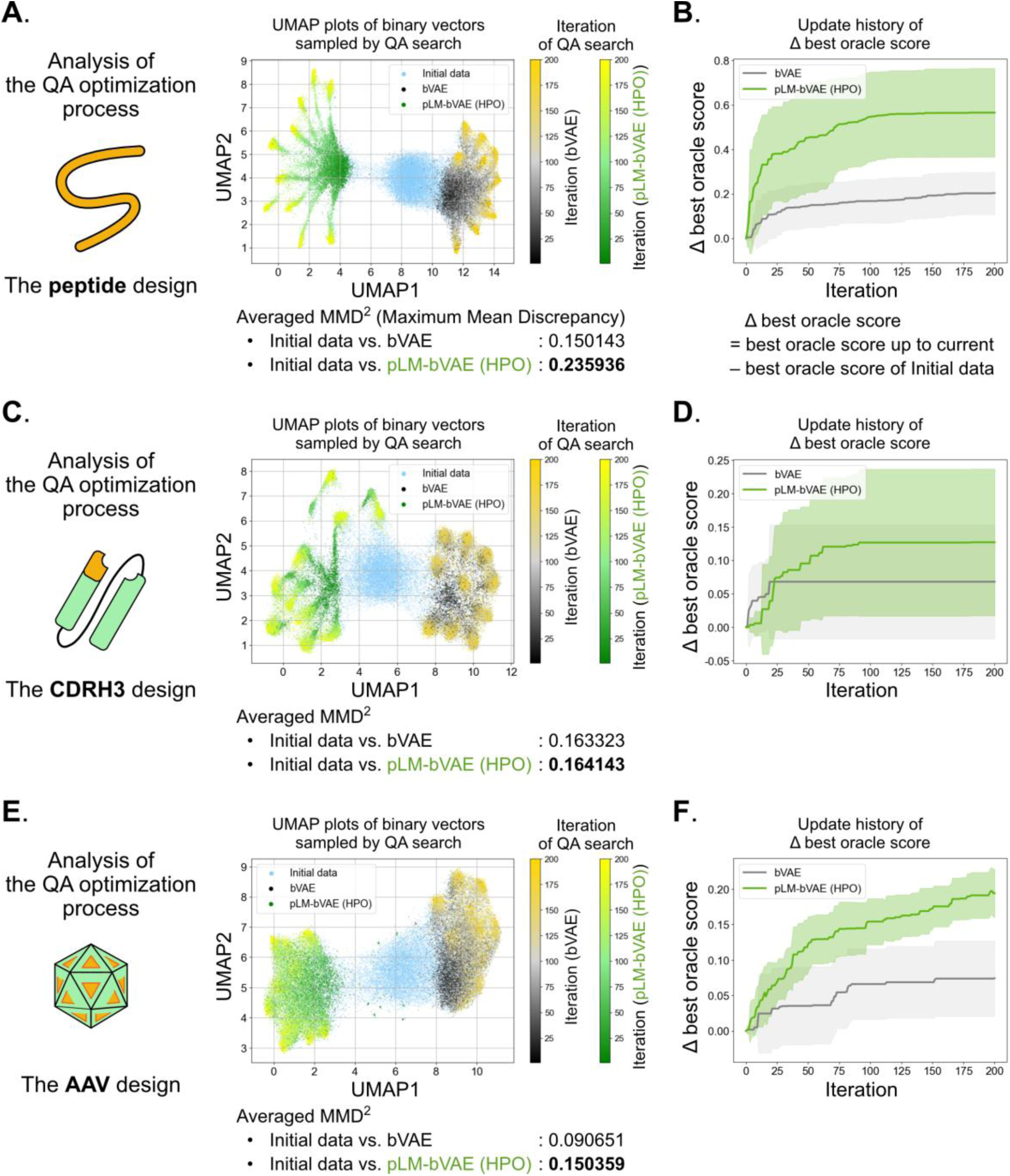
QA optimization using pLM-bVAE (HPO) showed a more directed search pattern with improved optimization efficiency. **(A, C, E)** UMAP plots of binary vectors of initial data and binary vectors sampled by QA search using bVAE/pLM-bVAE (HPO) on the peptide design **(A)**, the CDRH3 design **(C)**, and the AAV design **(E)**. The color bars indicate the iteration number of the QA searches. **(B, D, F)** Update history of Δ best oracle score by QA search using bVAE/pLM-bVAE (HPO) for each iteration on the peptide design **(B)**, the CDRH3 design **(D)**, and AAV design **(F)**. Δ best oracle score is calculated by subtracting the best oracle score of initial data from the best oracle score up to each iteration.

To examine whether this more directed search behavior was associated with improved QA optimization efficiency, we traced changes in Δ best oracle scores, calculated by subtracting the best oracle score of the initial data from the best oracle score obtained up to each iteration (Fig. 5B, 5D, 5F). Consistent with the larger MMD^2^ values, greater increases in Δ best oracle scores were observed in the QA optimization using pLM-bVAE, compared with that using bVAE, across all datasets. These results indicate that the greater distributional shift observed with pLM-bVAE (HPO) was associated with more effective improvement of the objective score during the QA optimization.

Taken together, our results suggest that incorporating pLM-bVAE (HPO) leads to a more directed search process in the binary latent space, which is associated with improved QA optimization efficiency. This improvement may facilitate QA-based protein sequence design.

## Author Contributions

H.H.U. and Y.S. conceived the method. H.H.U. wrote the code and performed the computational experiments. H.H.U. and Y.S. wrote the paper. A.T., K.T., M.U. and Y.S. helped model implementation and participated in the data interpretation. All authors read and approved the final version of the manuscript.

Conflict of interest: The authors declare that they have no competing interests.

## Supplementary data

Supplementary data is provided as a separate file.

## Acknowledgements

The computations were partially performed on the HOKUSHIN supercomputer at Department of Data Science, Kitasato University and the D-Wave Advantage system via the D-Wave Leap cloud platform.

## Data availability

pLM-bVAE software is available at https://github.com/hueda8/pLM-bVAE.

