## Supplementary Information for "pLM-bVAE: protein language model-enhanced binary representation improves quantum annealing-based protein sequence design"

### Supplementary text

**The architecture of bVAE.**

bVAE is a recurrent encoder-decoder model using gated recurrent unit (GRU)-based RNN [1] (Fig. S1). First, following the previous work using bVAE for binary encoding of amino acid sequences [2, 3], amino acids of the variant region were converted to 10 dimensional feature vectors using an amino acid descriptor. The feature vectors of the variant region were input to the bidirectional RNN-based bVAE encoder, producing probability vectors $\boldsymbol{p}$ through an element-wise sigmoid function. To obtain the binary latent vectors $\boldsymbol{z}$ from the probability vectors $\boldsymbol{p}$, straight-through estimator (STE) was employed [4]. During the forward pass, each element of the probability vectors $\boldsymbol{p}$ was binarized by applying the Heaviside step function and sampling $\boldsymbol{\xi}$ from an uniform distribution between 0 and 1, yielding the binary latent vectors $\boldsymbol{z}$:

$$z_i = step(p_i - \xi_i), \quad \xi_i \sim U(0,1)$$

where $i$ indexes the elements of the vector with 64 dimensions, $step(\cdot)$ denotes the Heaviside step function, and $U(0,1)$ denotes the uniform distribution. During the backward pass, STE defined the gradients through the step function were as follows:

$$\frac{\partial \mathcal{L}}{\partial p_i} := \frac{\partial \mathcal{L}}{\partial z_i}$$

where $\mathcal{L}$ denotes the loss function value.

To decode amino acid sequences from the binary latent vectors, the binary vectors were projected to the initial hidden state of the RNN-based bVAE decoder. At each decoding step $t$, the decoder produced a categorical probability distribution over amino acid tokens through a softmax function. During the bVAE training, the reconstruction loss was defined as the negative log-likelihood (NLL) of the ground-truth sequences:

$$\mathcal{L}_{NLL\ of\ bVAE} = -\frac{1}{B}\sum_{b=1}^{B}\sum_{t=1}^{L_b+1}\log \boldsymbol{p}_{\boldsymbol{t}}^{(b)}\left(y_t^{(b)}\right)$$

where $B$ denotes the mini-batch size, $L_b$ denotes the length of the $b$-th amino acid sequence, and $\boldsymbol{p}_{\boldsymbol{t}}^{(b)}$ and $y_t^{(b)}$ respectively denote the predicted probability distribution and its ground-truth token at the position $t$. During the inference, the most likely amino acid was selected from the probability distribution at each decoding step. Sequence generation was terminated when the <eos> token was generated.

**MOQA framework.**

First, to set the initial space searched by QA, 2,000 binary vectors were randomly sampled (Fig. S2). These binary vectors were reconstructed to 2,000 amino acid sequences using the trained bVAE/pLM-bVAE (HPO) decoders. The objective function values of these sequences were predicted by the three trained BiLSTM predictors. The predicted scores were averaged to be used as a computational oracle score. Then, the sequences were ranked using non-dominated sorting based on the three predicted scores. The ranks were used to define the function of MOQA score as follows:

$$y_{moqa} = \begin{cases} r/20 & (r \leq 20) \\ \log(1+r) & (r > 20) \end{cases}$$

where $y_{moqa}$ is the value of the MOQA score function, and $r$ is the rank of the sequence. Until this, the initial dataset that consists of 2,000 pairs of randomly sampled binary vectors and $y_{moqa}$ values was obtained. Since FM was the surrogate model for the MOQA score function, FM was trained on the initial dataset with the MSE loss function:

$$\mathcal{L}_{MSE} = \frac{1}{B}\sum_{b=1}^{B}\left|y_{moqa}{}^{(b)} - \widehat{y^{(b)}}\right|^2$$

where $B$ denotes the mini-batch size, which was set to 64, $y_{moqa}{}^{(b)}$ is the value of the MOQA score function of the $b$-th data, and $\widehat{y^{(b)}}$ is its prediction value by the FM. The trained FM was converted to QUBO, and QA was performed. QA sampled 10 reads as 64-dimensional binary vectors at an iteration with the annealing time of 20 μs. The sampled binary vectors were added to the existing dataset, and the FM was retrained using this additional dataset. This process was repeated for 200 iterations, generating 2,000 sequences. This QA-based sequence design procedure was repeated 10 times using different initial datasets, yielding approximately 20,000 sequences in total.

**Calculation of MMD with RBF kernel.**

Let $X = \{\boldsymbol{x}_1, \boldsymbol{x}_2, \cdots \boldsymbol{x}_m\}$ and $Y = \{\boldsymbol{y}_1, \boldsymbol{y}_2, \cdots \boldsymbol{y}_n\}$ denote the comparison distributions of binary vectors $\boldsymbol{x}_i$ and $\boldsymbol{y}_i$, respectively. $\mathrm{MMD}^2$ is given by:

$$\mathrm{MMD}^2(X,Y) = \frac{1}{m^2}\sum_{i=1}^{m}\sum_{j=1}^{m} k(\boldsymbol{x}_i, \boldsymbol{x}_j) + \frac{1}{n^2}\sum_{i=1}^{n}\sum_{j=1}^{n} k(\boldsymbol{y}_i, \boldsymbol{y}_j) - \frac{1}{mn}\sum_{i=1}^{m}\sum_{j=1}^{n} k(\boldsymbol{x}_i, \boldsymbol{y}_j)$$

where $k(\cdot,\cdot)$ is defined as RBF kernel similarity based on normalized Hamming distance of two binary vectors included in the two comparison distributions. Let $d_{Hamming}(\cdot,\cdot)$ denote normalized Hamming distance. The RBF kernel similarity is calculated as follows:

$$k(\cdot,\cdot) = exp\{-\gamma * d_{Hamming}(\cdot,\cdot)\}$$

where $\gamma$ is a hyperparameter of RBF kernel. In this study, the value of $\gamma$ was determined by median heuristic method [5]. Let $d_{median}$ denote the median normalized Hamming distance between the binary vectors of the initial data and sampled by the QA searches using bVAE or pLM-bVAE (HPO). Then, $\gamma$ is calculated as follows:

$$\gamma = \frac{1}{d_{median}}$$

**Training and hyperparameter optimization of BiLSTM predictors for objective functions**

Bayesian optimization of the hyperparameters was performed with 5-fold cross-validation of the dataset containing the training and validation sets using PARROT package [6]. First, amino acids in the variant region were converted to 3 dimensional feature vectors using an amino acid descriptor, Z-scales (3), which has high modeling capability [7, 8]. Then, to optimize three hyperparameters (i.e., the learning rate, the number of layers, and the hidden size), Bayesian optimization was performed with 100 epochs. The three hyperparameter sets with the lowest validation MAE values were selected. Then, three BiLSTM predictors were trained with the three hyperparameter sets, respectively (Fig. S8A). The trainings were stopped early using the validation set.

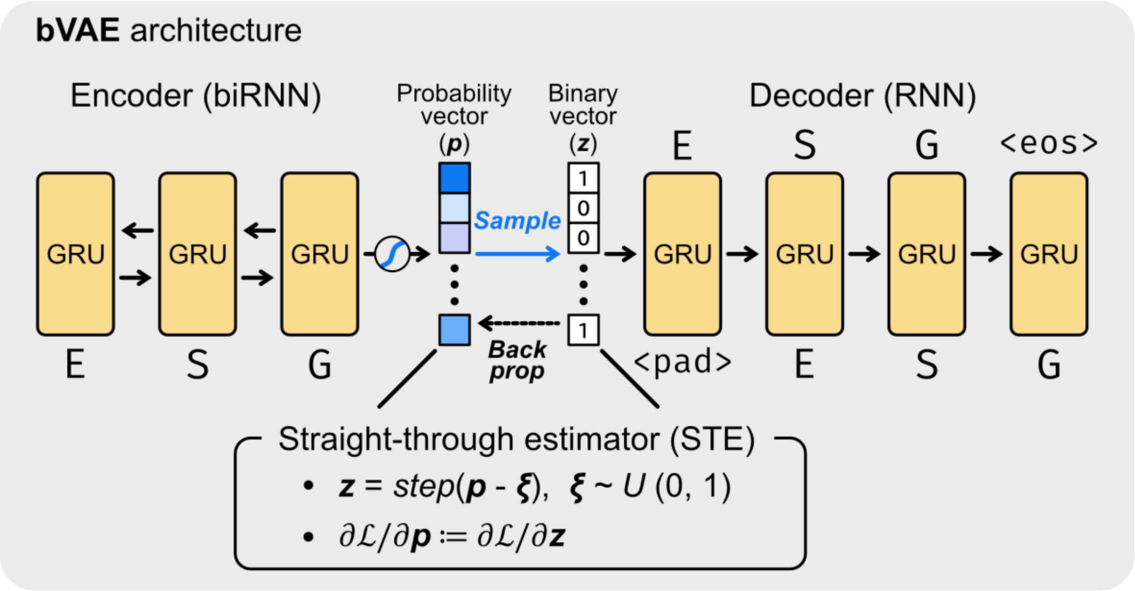


**Figure S1. The architecture of bVAE.**

bVAE is a GRU-based recurrent encoder-decoder model. The bidirectional RNN-based encoder receives an amino acid sequence using an amino acid descriptor or a pLM embedding, which produces a probability vector $\boldsymbol{p}$ through an element-wise sigmoid function. To obtain a binary latent vector $\boldsymbol{z}$, STE is employed. The each element of the probability vector $\boldsymbol{p}$ is binarized by applying the Heaviside step function and sampling $\xi$ from the uniform distribution $U(0, 1)$, yielding the binary latent vector $\boldsymbol{z}$. During the backward pass, STE defined the gradient through the step function as $\partial\mathcal{L}/\partial p_i \coloneqq \partial\mathcal{L}/\partial z_i$. During the inference, the RNN-based decoder receives the binary latent vector, generating amino acid sequences. Sequence generation was terminated when the <eos> token was generated.

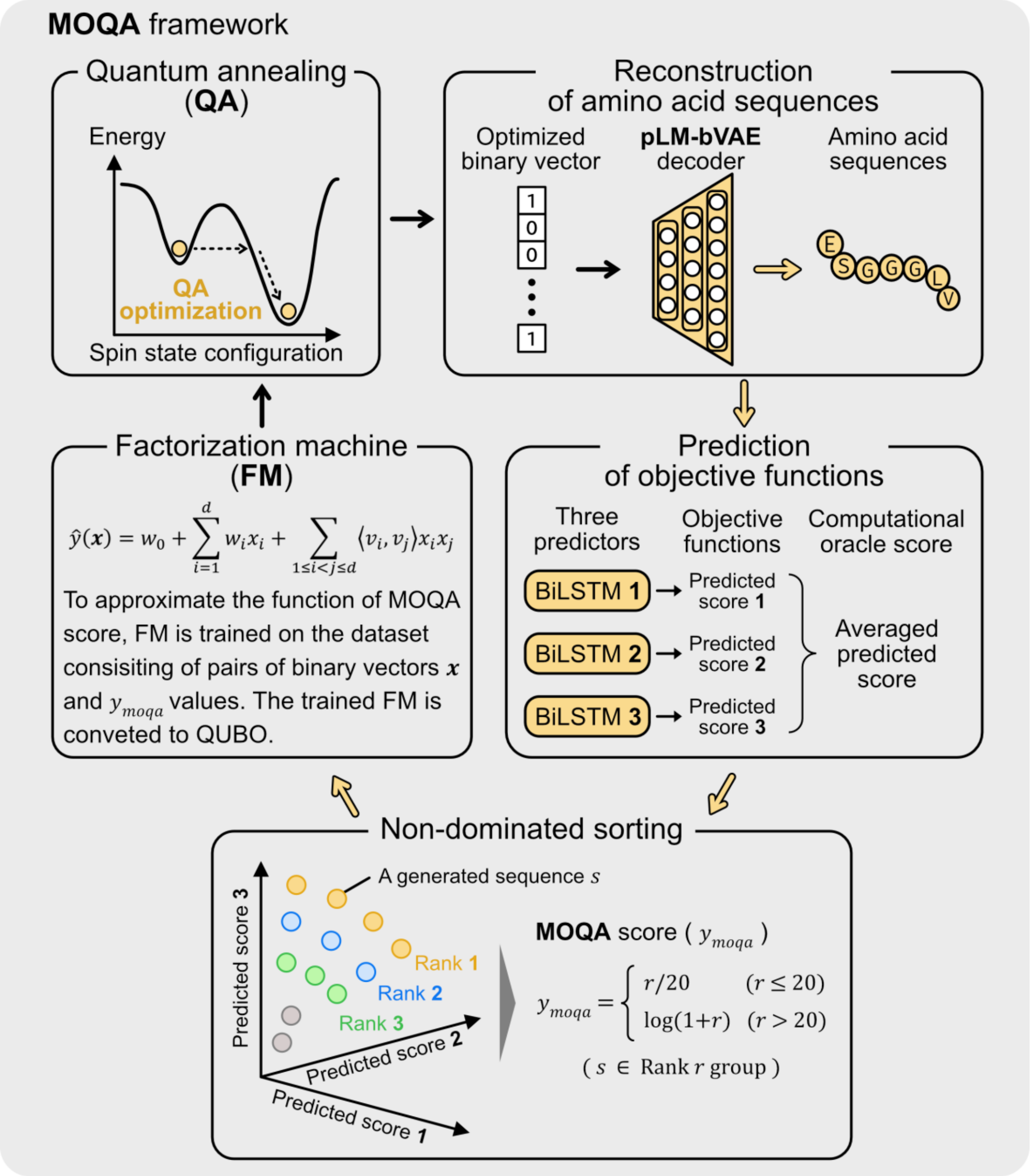


**Figure S2. Multi-objective optimization by quantum annealing (MOQA) framework.**

To generate sequences with a high objective value, MOQA framework was employed. In this study, multi objectives were the same objective for each protein design, such as $IC_{50}$ for the peptide design, $K_D$ for the CDRH3 design, and the viability for the AAV design. First, randomly sampled binary vectors (not optimized) were reconstructed to amino acid sequences using a trained pLM-bVAE decoder. The objective function values of the amino acid sequences were predicted by three trained BiLSTM predictors. These predicted scores were averaged to be used as a computational oracle score. Then, the sequences were ranked using non-dominated sorting based on the three predicted scores. The ranks were used to calculate a MOQA score ($y_{moqa}$). Until this, the dataset that consists of pairs of randomly sampled binary vectors and $y_{moqa}$ values was obtained. To approximate the function of MOQA score, FM was trained on the dataset with the MSE loss, because FM was employed as surrogate model for the function of MOQA score. The trained FM was converted to QUBO that can be solved by QA. QA optimization sampled optimized binary vectors. The amino acid sequences with high objective values were obtained by reconstructing them from the QA-optimized binary vectors using the pLM-bVAE decoder. The optimized binary vectors were added to the existing datasets, and the FM was retrained using this additional dataset. This process was repeated for 200 iterations.

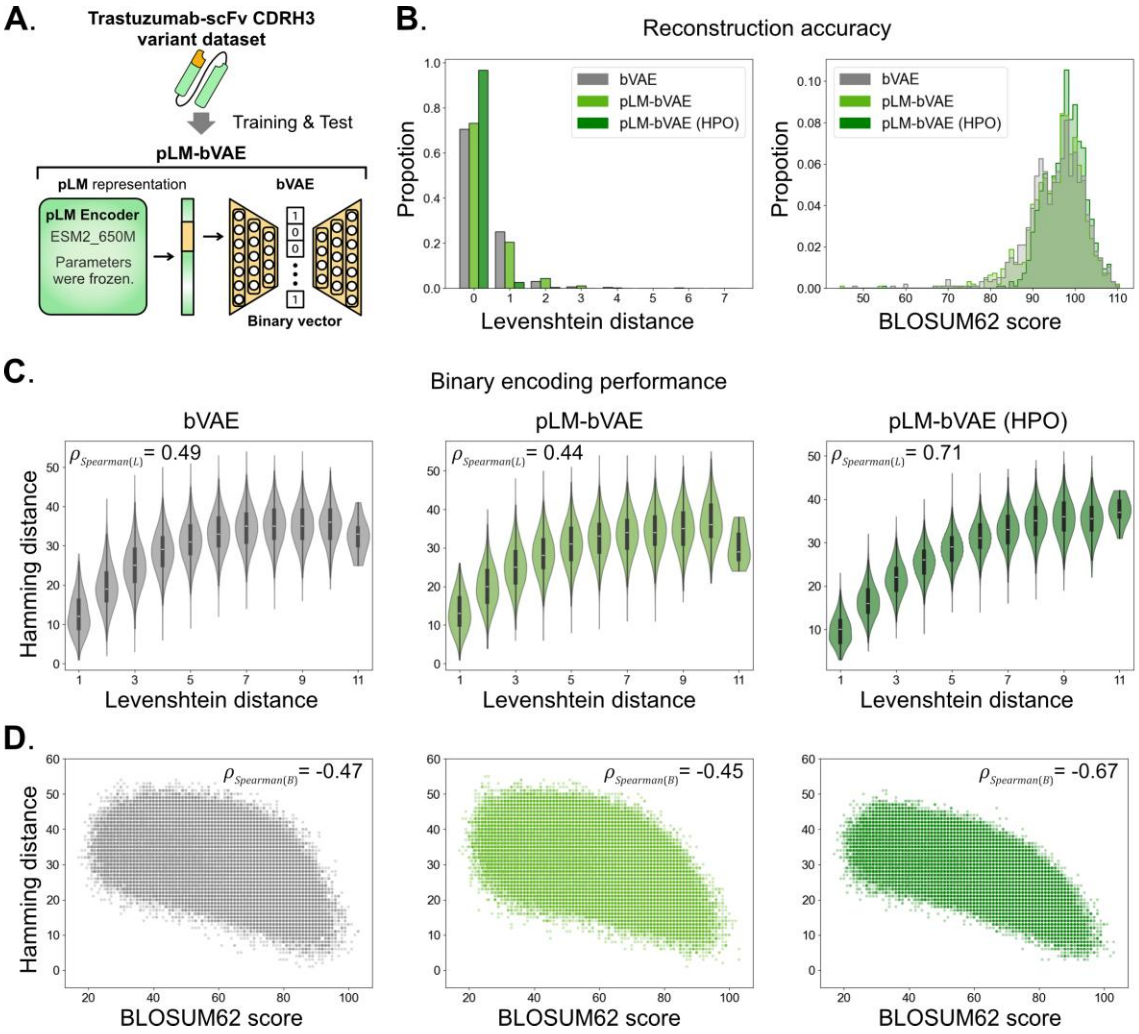


**Figure S3. pLM-bVAE showed higher reconstruction accuracy and binary encoding performance than bVAE on the Trastuzumab-scFv CDRH3 variant dataset.**

**(A)** To train and test bVAE and pLM-bVAE, the Trastuzumab-scFv CDRH3 variant dataset was used. pLM-bVAE was composed of ESM2 650M encoder with the model parameters frozen and bVAE.

**(B)** Reconstruction accuracy of the trained bVAE, pLM-bVAE, and pLM-bVAE (HPO) on the test data. **Left**: Distributions of Levenshtein distance between input sequences and output sequences through the models. **Right**: Distributions of BLOSUM62 score between input sequences and output sequences through the models.

**(C, D)** Binary encoding performance of the trained bVAE (left), pLM-bVAE (middle), and pLM-bVAE (HPO) (right) on the test data. **(C)** Correlation between pairwise Levenshtein distance of the test sequences and pairwise Hamming distance of the binary vectors encoded from their sequences. $\rho_{Spearman(L)}$ shows the Spearman correlation coefficient value. **(D)** Correlation between pairwise BLOSUM62 score of the test sequences and pairwise Hamming distance of the binary vectors encoded from their sequences. $\rho_{Spearman(B)}$ shows the Spearman correlation coefficient value.

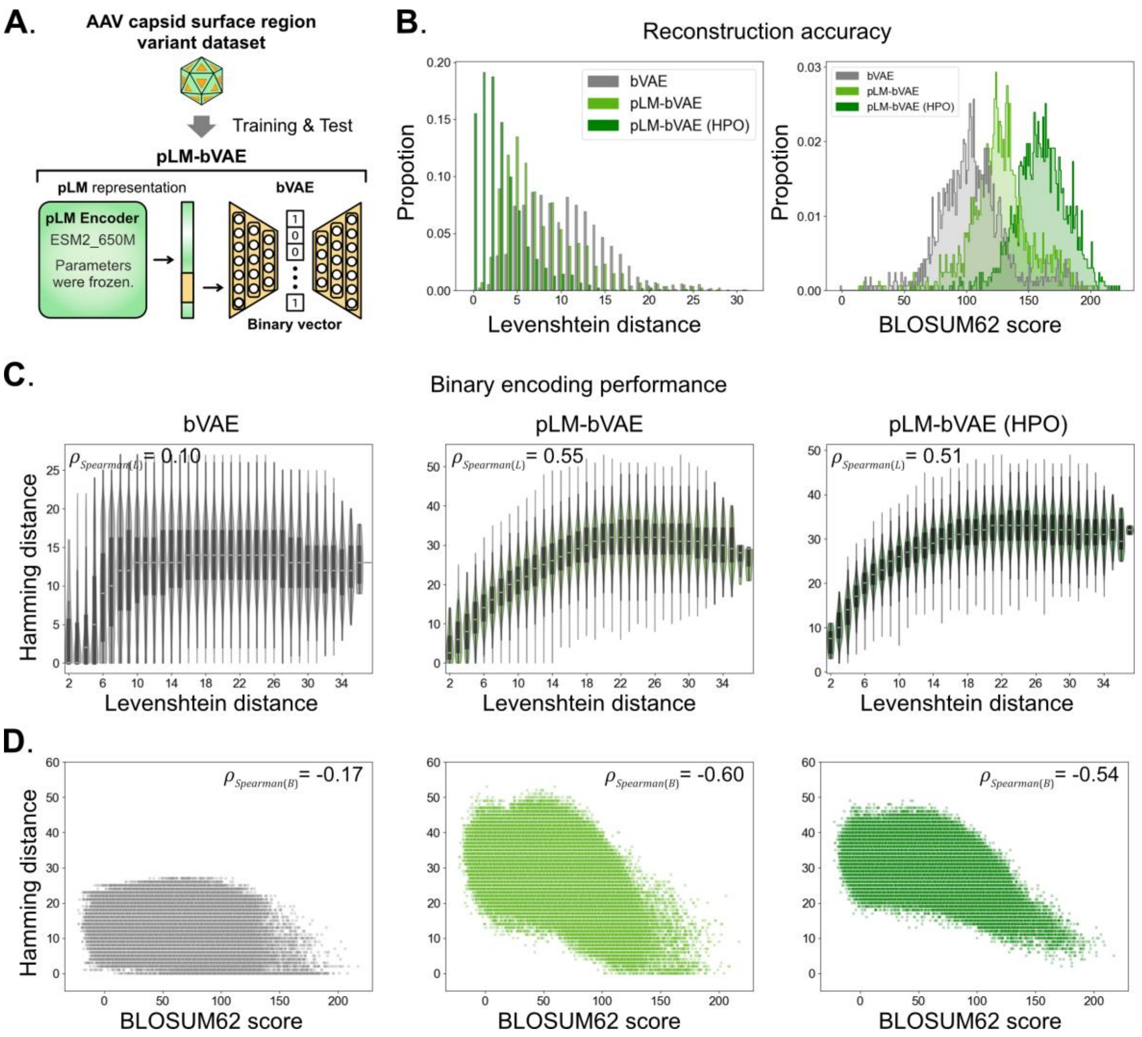


**Figure S4. pLM-bVAE showed higher reconstruction accuracy and binary encoding performance than bVAE on the AAV capsid surface region variant dataset.**

**(A)** To train and test bVAE and pLM-bVAE, the AAV capsid surface region variant dataset was used. pLM-bVAE was composed of ESM2 650M encoder with the model parameters frozen and bVAE.

**(B)** Reconstruction accuracy of the trained bVAE, pLM-bVAE, and pLM-bVAE (HPO) on the test data. **Left**: Distributions of Levenshtein distance between input sequences and output sequences through the models. **Right**: Distributions of BLOSUM62 score between input sequences and output sequences through the models.

**(C, D)** Binary encoding performance of the trained bVAE (left), pLM-bVAE (middle), and pLM-bVAE (HPO) (right) on the test data. **(C)** Correlation between pairwise Levenshtein distance of the test sequences and pairwise Hamming distance of the binary vectors encoded from their sequences. $\rho_{Spearman(L)}$ shows the Spearman correlation coefficient value. **(D)** Correlation between pairwise BLOSUM62 score of the test sequences and pairwise Hamming distance of the binary vectors encoded from their sequences. $\rho_{Spearman(B)}$ shows the Spearman correlation coefficient value.

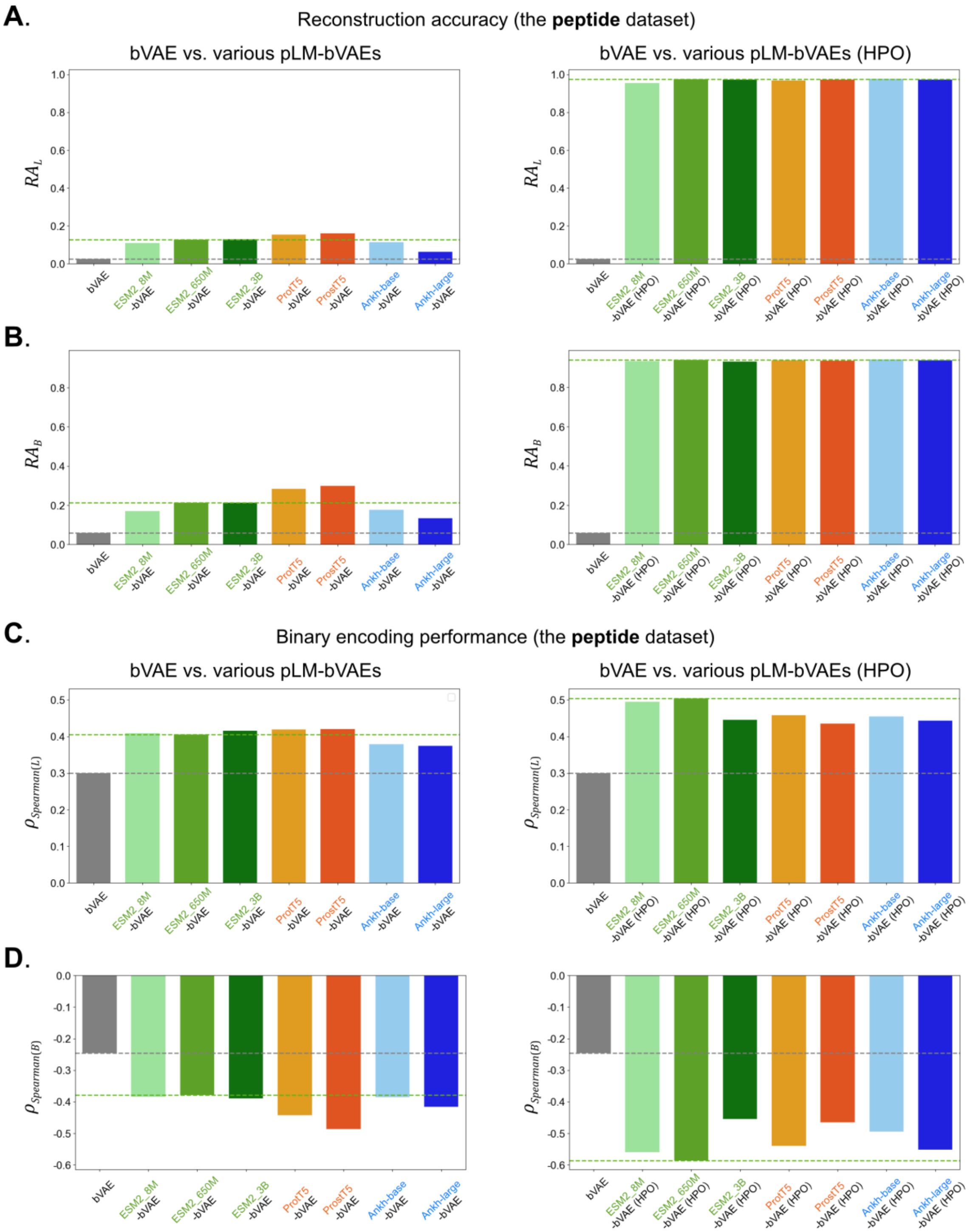


**Figure S5. The comparison of performances of bVAE and various pLM-bVAEs on the MHC-I binding peptide dataset.**

**(A, B)** Reconstruction accuracy of the trained bVAE vs. the trained various pLM-bVAEs (left) and the trained bVAE vs. the trained various pLM-bVAEs (HPO) (right) on the test data. **(A)** $RA_L$ shows the ratio of sequences with Levenshtein distance between input sequences and output sequences of 0. **(B)** $RA_B$ shows the ratio of sequences with BLOSUM62 score between input sequences and output sequences higher than the 95th percentile BLOSUM62 score of bVAE. The horizontal gray and green dashed lines show the performance scores of bVAE and ESM2_650M-bVAE (HPO), respectively. The types of encoders are listed in Table 2.

**(C, D)** Binary encoding performance of the trained bVAE vs. the trained various pLM-bVAEs (left) and the trained bVAE vs. the trained various pLM-bVAEs (HPO) (right) on the test data. **(C)** $\rho_{Spearman(L)}$ shows the Spearman correlation coefficient value between pairwise Levenshtein distance of the test sequences and pairwise Hamming distance of the binary vectors encoded from their sequences. **(D)** $\rho_{Spearman(L)}$ shows the Spearman correlation coefficient value between pairwise BLOSUM62 score of the test sequences and pairwise Hamming distance of the binary vectors encoded from their sequences.

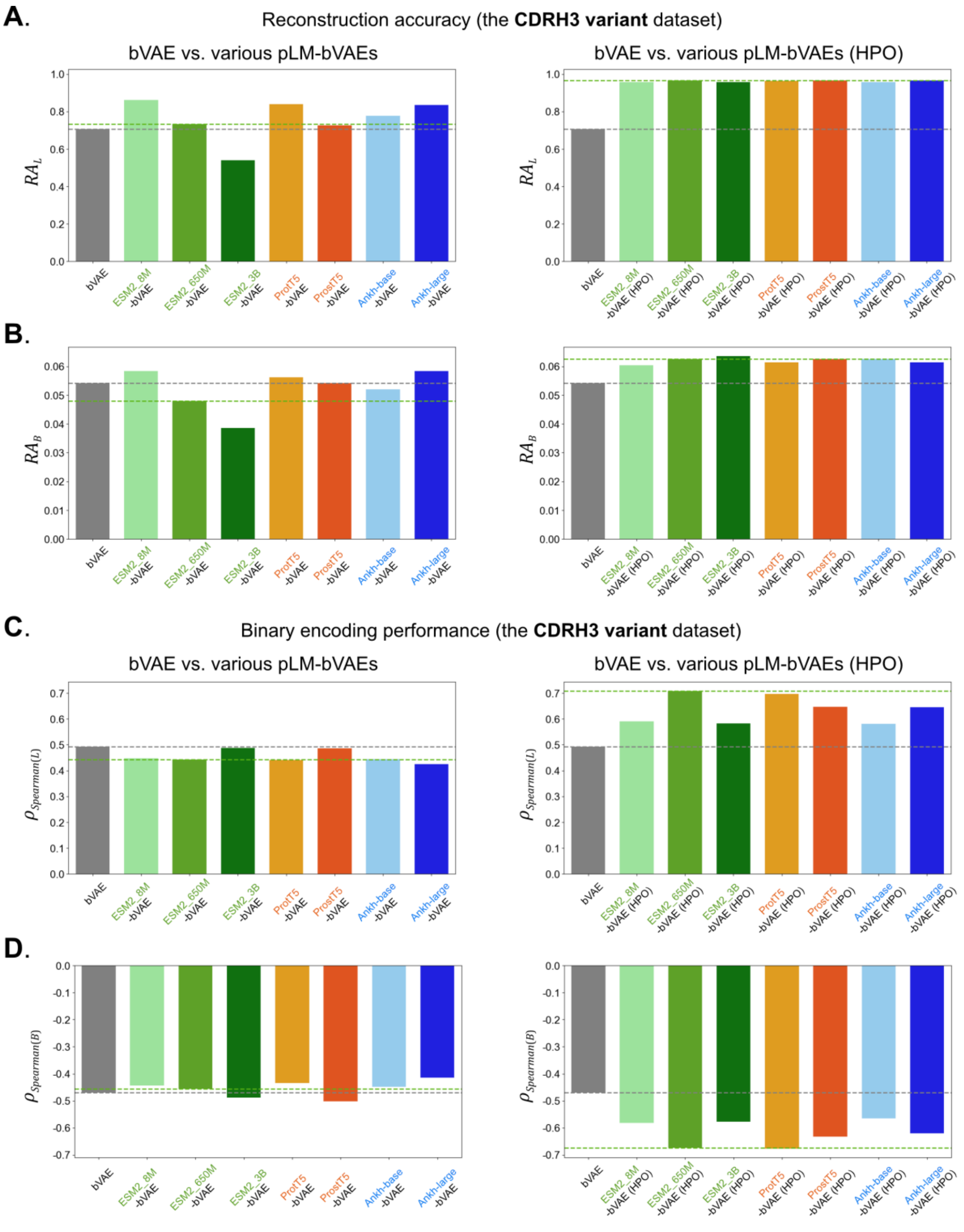


**Figure S6. The comparison of performance of bVAE and various pLM-bVAEs on the Trastuzumab-scFv CDRH3 variants dataset.**

**(A, B)** Reconstruction accuracy of the trained bVAE vs. the trained various pLM-bVAEs (left) and the trained bVAE vs. the trained various pLM-bVAEs (HPO) (right) on the test data. **(A)** $RA_L$ shows the ratio of sequences with Levenshtein distance between input sequences and output sequences of 0. **(B)** $RA_B$ shows the ratio of sequences with BLOSUM62 score between input sequences and output sequences higher than the 95th percentile BLOSUM62 score of bVAE. The horizontal gray and green dashed lines show the performance scores of bVAE and ESM2_650M-bVAE (HPO), respectively.

**(C, D)** Binary encoding performance of the trained bVAE vs. the trained various pLM-bVAEs (left) and the trained bVAE vs. the trained various pLM-bVAEs (HPO) (right) on the test data. **(C)** $\rho_{Spearman(L)}$ shows the Spearman correlation coefficient value between pairwise Levenshtein distance of the test sequences and pairwise Hamming distance of the binary vectors encoded from their sequences. **(D)** $\rho_{Spearman(L)}$ shows the Spearman correlation coefficient value between pairwise BLOSUM62 score of the test sequences and pairwise Hamming distance of the binary vectors encoded from their sequences.

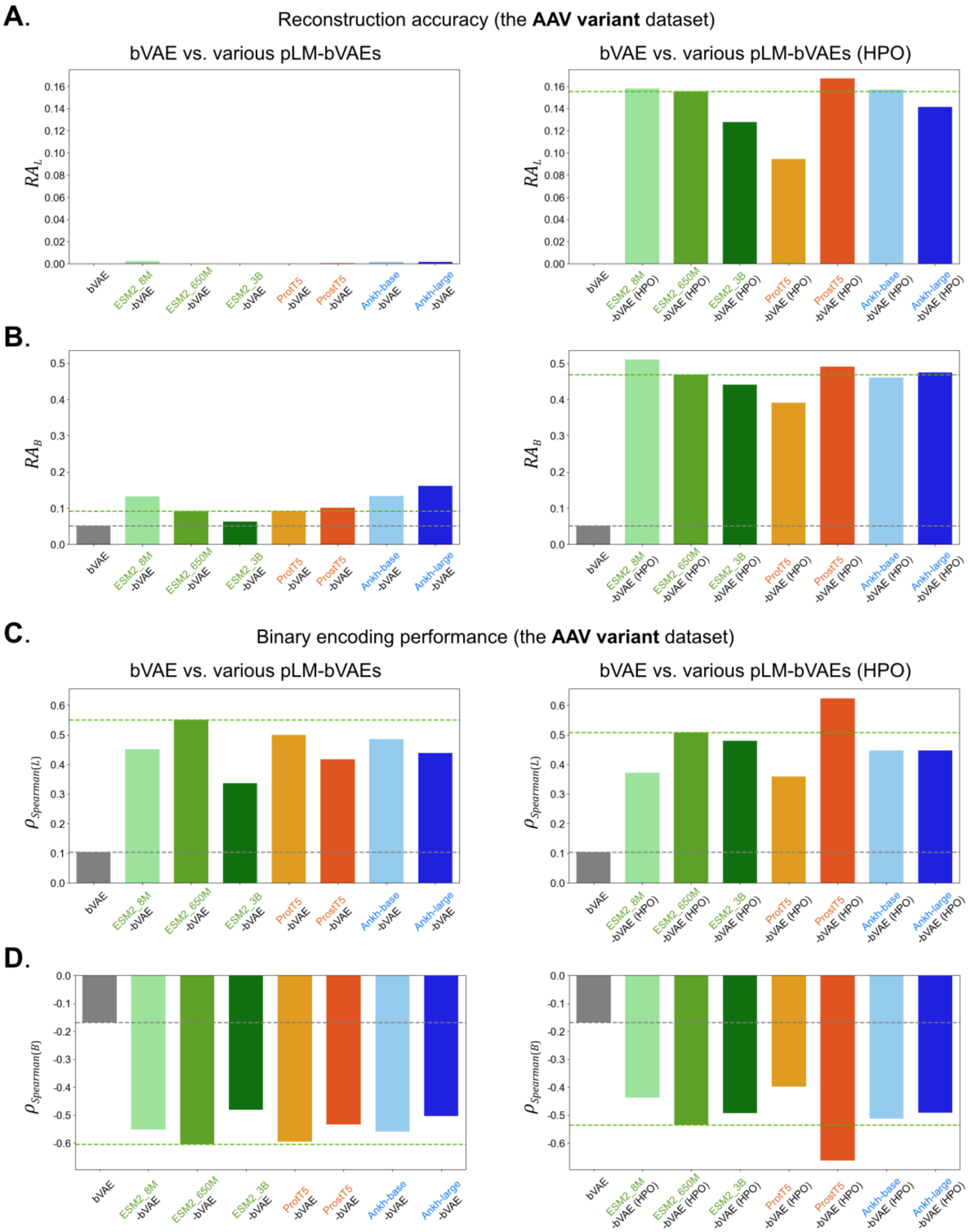


**Figure S7. The comparison of performance of bVAE and various pLM-bVAEs on the AAV capsid surface region variant dataset.**

**(A, B)** Reconstruction accuracy of the trained bVAE vs. the trained various pLM-bVAEs (left) and the trained bVAE vs. the trained various pLM-bVAEs (HPO) (right) on the test data. **(A)** $RA_L$ shows the ratio of sequences with Levenshtein distance between input sequences and output sequences of 0. **(B)** $RA_B$ shows the ratio of sequences with BLOSUM62 score between input sequences and output sequences higher than the 95th percentile BLOSUM62 score of bVAE. The horizontal gray and green dashed lines show the performance scores of bVAE and ESM2_650M-bVAE (HPO), respectively.

**(C, D)** Binary encoding performance of the trained bVAE vs. the trained various pLM-bVAEs (left) and the trained bVAE vs. the trained various pLM-bVAEs (HPO) (right) on the test data. **(C)** $\rho_{Spearman(L)}$ shows the Spearman correlation coefficient value between pairwise Levenshtein distance of the test sequences and pairwise Hamming distance of the binary vectors encoded from their sequences. **(D)** $\rho_{Spearman(L)}$ shows the Spearman correlation coefficient value between pairwise BLOSUM62 score of the test sequences and pairwise Hamming distance of the binary vectors encoded from their sequences.

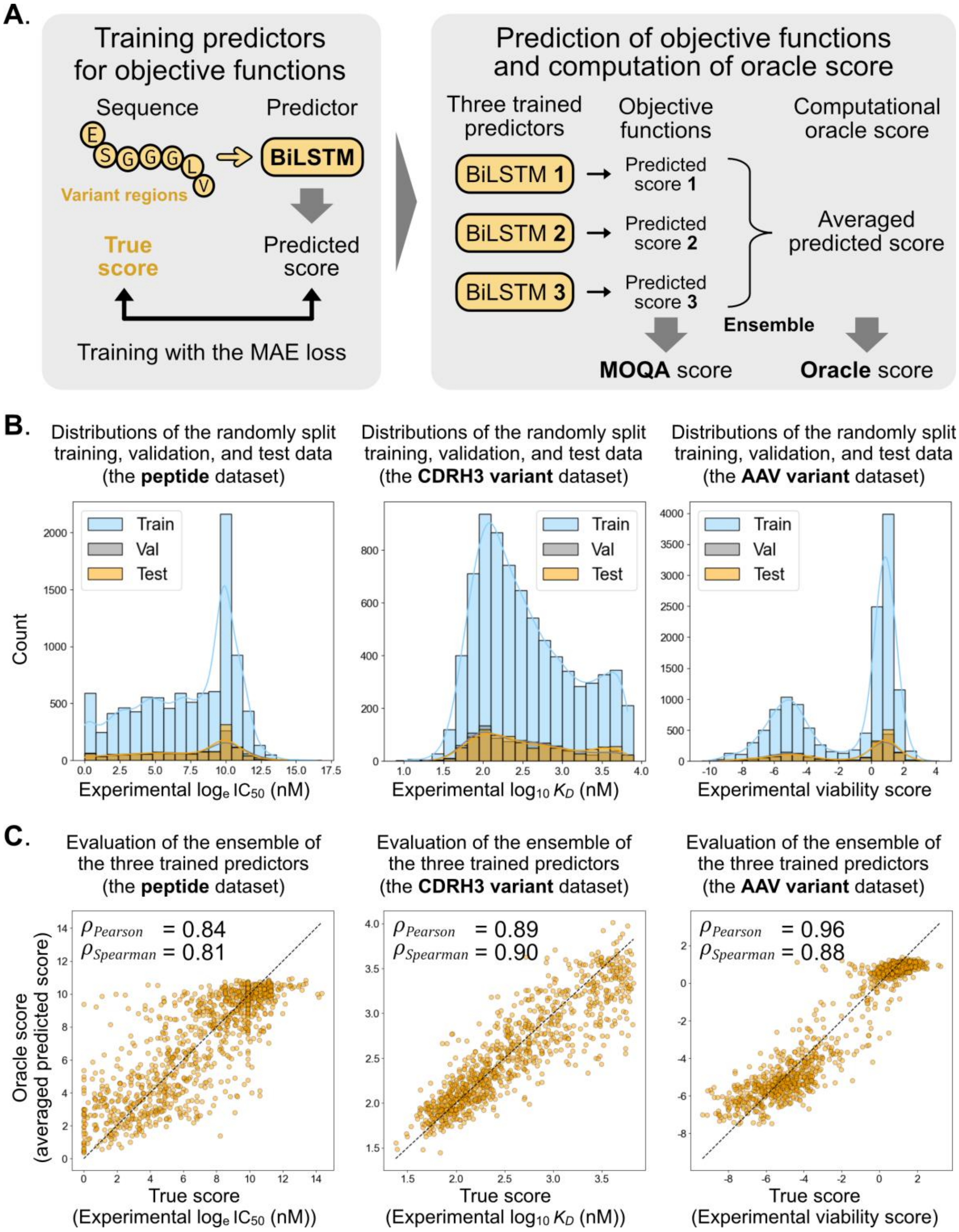


226

227 **Figure S8. Training three predictors for objective functions on each protein variant**

228 **dataset and evaluation of the ensemble of the three trained predictors on the test**

229 **data.**

**(A)** Training BiLSTM predictors with Bayesian optimized hyperparameters using the MAE loss. The three trained predictors output predicted scores as objective function values, which were used for non-dominated sorting of MOQA (Fig. S2). The ensemble of the three trained predictor output an average of predicted score as a computational oracle score for the evaluation of QA-based protein sequence design performance.
**(B)** Distributions of the randomly split training, validation, and test data of the peptide dataset (left), the CDRH3 variant dataset (middle), and the AAV variant dataset (right).
**(C)** Evaluation of the ensemble of the three trained predictor on the test data of the peptide dataset (left), the CDRH3 variant dataset (middle), and the AAV variant dataset (right). $\rho_{Pearson}$ and $\rho_{Spearman}$ respectively show the Pearson correlation coefficient value and the Spearman correlation value between the experimentally measured scores and the averaged predicted scores.

| Model | $RA_L$ | $RA_B$ | $\rho_{Spearman(L)}$ | $\rho_{Spearman(B)}$ | $S_{total}$ |
|---|---|---|---|---|---|
| bVAE | 0.024671 | 0.057566 | 0.300216 | -0.245869 | 0.000000 |
| ESM2_8M-bVAE | 0.108553 | 0.169408 | 0.408899 | -0.384179 | 0.178571 |
| ESM2_650M-bVAE | 0.126645 | 0.212993 | 0.405072 | -0.378793 | 0.223214 |
| ESM2_3B-bVAE | 0.129934 | 0.212993 | 0.415990 | -0.389251 | 0.330357 |
| ProtT5-bVAE | 0.152961 | 0.282895 | 0.419199 | -0.443401 | 0.428571 |
| ProstT5-bVAE | 0.161184 | 0.298520 | 0.419849 | -0.487101 | 0.535714 |
| Ankh-base-bVAE | 0.112664 | 0.175987 | 0.379267 | -0.386071 | 0.196429 |
| Ankh-large-bVAE | 0.062500 | 0.134046 | 0.374223 | -0.416721 | 0.142857 |
| ESM2_8M-bVAE (HPO) | 0.954770 | 0.934211 | 0.494805 | -0.559671 | 0.767857 |
| ESM2_650M-bVAE (HPO) | 0.974507 | 0.939145 | **0.503360** | **-0.586203** | **0.964286** |
| ESM2_3B-bVAE (HPO) | 0.972862 | 0.930921 | 0.445036 | -0.454839 | 0.651786 |
| ProtT5-bVAE (HPO) | 0.967928 | 0.937500 | 0.458436 | -0.540027 | 0.776786 |
| ProstT5-bVAE (HPO) | 0.972862 | 0.935855 | 0.434962 | -0.465506 | 0.669643 |
| Ankh-base-bVAE (HPO) | **0.976151** | **0.941612** | 0.454249 | -0.494847 | 0.875000 |
| Ankh-large-bVAE (HPO) | 0.972039 | 0.937500 | 0.443232 | -0.552195 | 0.758929 |

**Table S1. The performance metrics of the trained bVAE and various pLM-bVAEs on the MHC-I binding peptide dataset.**

| Model | $RA_L$ | $RA_B$ | $\rho_{Spearman(L)}$ | $\rho_{Spearman(B)}$ | $S_{total}$ |
|---|---|---|---|---|---|
| bVAE | 0.706250 | 0.054167 | 0.493083 | -0.470401 | 0.294643 |
| ESM2_8M-bVAE | 0.862500 | 0.058333 | 0.447523 | -0.444021 | 0.348214 |
| ESM2_650M-bVAE | 0.732292 | 0.047917 | 0.442781 | -0.455279 | 0.178571 |
| ESM2_3B-bVAE | 0.540625 | 0.038542 | 0.487555 | -0.488711 | 0.214286 |
| ProtT5-bVAE | 0.838542 | 0.056250 | 0.440795 | -0.434410 | 0.232143 |
| ProstT5-bVAE | 0.726042 | 0.054167 | 0.486897 | -0.502157 | 0.312500 |
| Ankh-base-bVAE | 0.776042 | 0.052083 | 0.443669 | -0.448954 | 0.214286 |
| Ankh-large-bVAE | 0.835417 | 0.058333 | 0.425321 | -0.414725 | 0.205357 |
| ESM2_8M-bVAE (HPO) | 0.956250 | 0.060417 | 0.590971 | -0.581298 | 0.651786 |
| ESM2_650M-bVAE (HPO) | 0.966632 | 0.062565 | **0.708118** | -0.674729 | **0.946429** |
| ESM2_3B-bVAE (HPO) | 0.956250 | **0.063542** | 0.582583 | -0.577955 | 0.723214 |
| ProtT5-bVAE (HPO) | 0.962500 | 0.061458 | 0.696108 | **-0.677695** | 0.848214 |
| ProstT5-bVAE (HPO) | **0.966667** | 0.062500 | 0.646769 | -0.633332 | 0.883929 |
| Ankh-base-bVAE (HPO) | 0.957292 | 0.062500 | 0.580385 | -0.565313 | 0.669643 |
| Ankh-large-bVAE (HPO) | 0.965625 | 0.061458 | 0.645413 | -0.620086 | 0.776786 |

**Table S2. The performance metrics of the trained bVAE and various pLM-bVAEs on the Trastuzumab-scFv CDRH3 variant dataset.**

| Model | $RA_L$ | $RA_B$ | $\rho_{Spearman(L)}$ | $\rho_{Spearman(B)}$ | $S_{total}$ |
|---|---|---|---|---|---|
| bVAE | 0.000000 | 0.050746 | 0.104388 | -0.167983 | 0.026786 |
| ESM2_8M-bVAE | 0.002388 | 0.132537 | 0.451944 | -0.552152 | 0.535714 |
| ESM2_650M-bVAE | 0.000000 | 0.091343 | 0.550175 | -0.604452 | 0.526786 |
| ESM2_3B-bVAE | 0.000000 | 0.062687 | 0.336090 | -0.480772 | 0.116071 |
| ProtT5-bVAE | 0.000000 | 0.091940 | 0.499368 | -0.595489 | 0.491071 |
| ProstT5-bVAE | 0.000597 | 0.101493 | 0.417377 | -0.533843 | 0.357143 |
| Ankh-base-bVAE | 0.001791 | 0.133731 | 0.485289 | -0.558679 | 0.580357 |
| Ankh-large-bVAE | 0.001791 | 0.161791 | 0.438962 | -0.503477 | 0.419643 |
| ESM2_8M-bVAE (HPO) | 0.158209 | **0.509851** | 0.371637 | -0.437447 | 0.571429 |
| ESM2_650M-bVAE (HPO) | 0.155224 | 0.468657 | 0.507832 | -0.535027 | 0.767857 |
| ESM2_3B-bVAE (HPO) | 0.127761 | 0.441194 | 0.479945 | -0.493485 | 0.571429 |
| ProtT5-bVAE (HPO) | 0.094328 | 0.391045 | 0.358794 | -0.397954 | 0.339286 |
| ProstT5-bVAE (HPO) | **0.167164** | 0.490149 | **0.623048** | **-0.662779** | **0.982143** |
| Ankh-base-bVAE (HPO) | 0.157015 | 0.460299 | 0.446974 | -0.512239 | 0.642857 |
| Ankh-large-bVAE (HPO) | 0.141493 | 0.474030 | 0.446927 | -0.490790 | 0.571429 |

**Table S3. The performance metrics of the trained bVAE and various pLM-bVAEs on the AAV capsid surface region variant dataset.**

[8] van Westen GJ, Swier RF, Cortes-Ciriano I, Wegner JK, Overington JP, IJzerman AP, van Vlijmen HW, Bender A. Benchmarking of protein descriptor

sets in proteochemometric modeling (part 2): modeling performance of 13 amino acid descriptor sets. Journal of Cheminformatics. 2013;5(1):42. doi:10.1186/1758-2946-5-42
